# A Tale of Twins: The Distinct Roles of Intracellular and Extracellular Formate in Regulating Flagellar and Pathogenicity Island-1 Genes in *Salmonella* Typhimurium

**DOI:** 10.1101/2024.01.30.577997

**Authors:** Debapriya Mukherjee, Dipshikha Chakravortty

**Affiliations:** Department of Microbiology and Cell Biology, Division of Biological Sciences, Indian Institute of Science, Bangalore, India; School of Biology, Indian Institute of Science Education and Research, Thiruvananthapuram, 695551

**Keywords:** Flagella, Formate, Intracellular pH, Membrane depolarization, Short Chain Fatty Acids (SCFAs)

## Abstract

Host-derived short-chain fatty acids (SCFAs) are extensively being studied for their role in the virulence and pathogenesis of *Salmonella* Typhimurium (STM). Formate, an SCFA present in the ileum, functions as a signalling molecule to enhance STM invasion. However, the role of intracellular formate in *Salmonella* virulence remains poorly understood. To investigate this, we generated knockouts of *pflB* (pyruvate-formate lyase) and *focA* (formate transporter). Disruption of formate production through *pflB* deletion led to reduced flagellation and increased *hilA* and *prgH* expression, attributed to elevated intracellular pH and membrane damage. This suppression of flagellar machinery drives a shift from adhesion to invasion, regulated by RpoE via the CsrA/*csrB* pathway. Additionally, we demonstrate that upon compensation for intracellular deficiency of formate, STM *ΔpflB* starts to utilize formate as a signalling molecule to regulate downstream processes. To the best of our knowledge, this study is the first to establish the critical role of the *pflB* gene in maintaining intracellular pH and controlling virulence gene expression in STM. Lastly, our findings emphasize the importance of modulating *pflB* expression across intestinal regions to optimize STM invasion.

**Graphical abstract:** 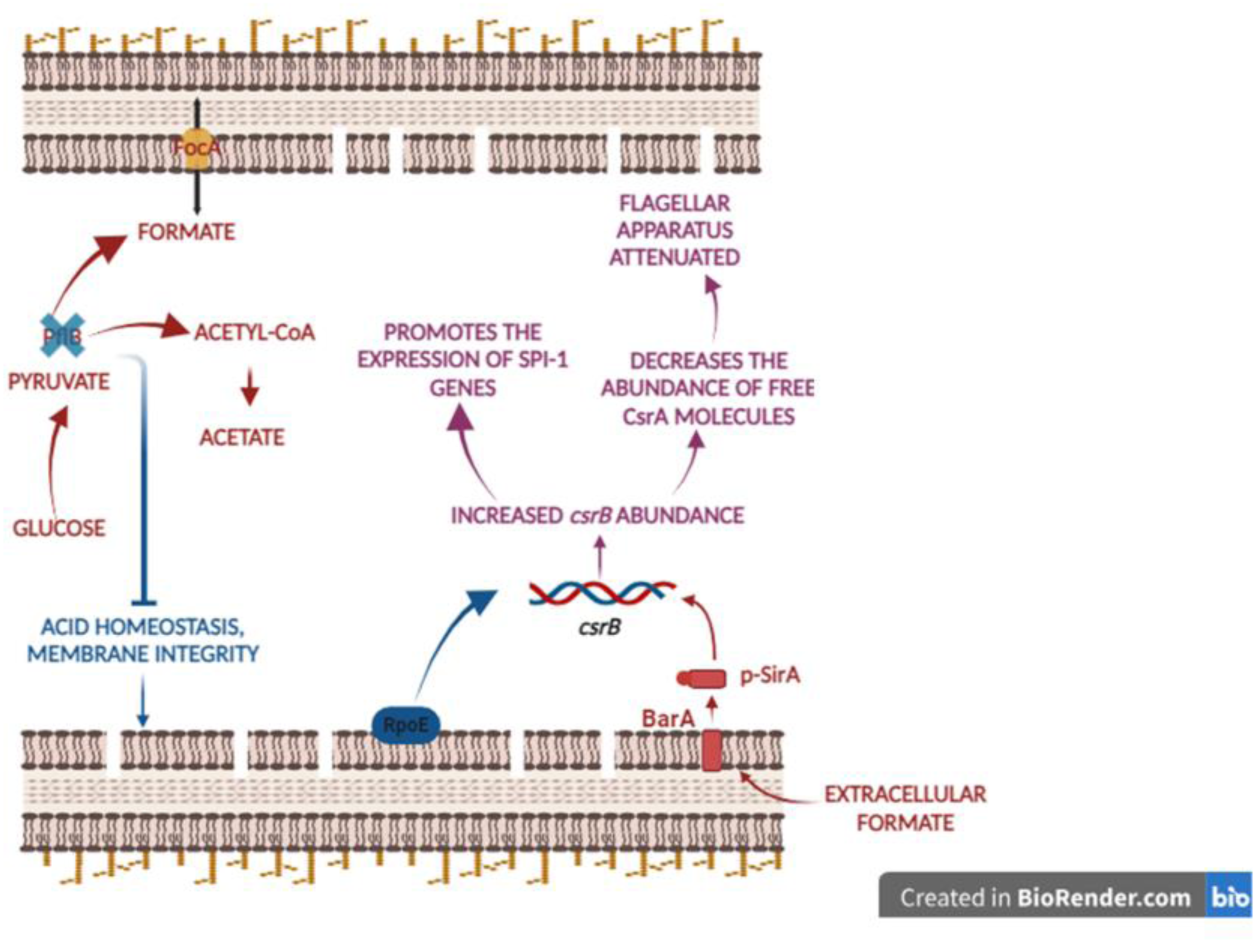

## Introduction

The genus *Salmonella* consists of Gram-negative gammaproteobacteria that infect a wide range of human and animal hosts [1]. Non-typhoidal serovars, such as *Salmonella* Typhimurium, are commonly associated with self-limiting diarrheal illness with relatively low fatality. According to the Global Burden of Diseases, Injuries, and Risk Factors Study (GBD) 2017, there were an estimated 95.1 million cases of *Salmonella* enterocolitis worldwide, resulting in 50,771 deaths [2-4]. Although non-typhoidal infections are typically confined to the intestines, these serovars can occasionally invade sterile tissues, leading to bacteraemia, meningitis, and other focal infections, which significantly increase mortality rates.[5].

*Salmonella* Typhimurium enters the host via contaminated food and water. One of the defence mechanisms in the host is the presence of short-chain fatty acids (SCFAs) in the small intestine. SCFAs are produced through the anaerobic fermentation of non-digestible polysaccharides, such as resistant starches and dietary fibres. The SCFAs found in the intestine—formate (C1), acetate (C2), propionate (C3), and butyrate (C4), with concentrations reaching up to 10–100 mM. Recent studies have extensively explored the role of SCFAs ((like butyrate and propionate) in *Salmonella* infections [6-10]. For example, butyrate downregulates the expression of pathogenicity island-1 (SPI-1) genes in *S.* Typhimurium, thereby reducing bacterial invasion and translocation from the intestines to the bloodstream [11-13].

Formate present in the intestine has been previously shown to act as a diffusible signal promoting *Salmonella* Typhimurium invasion. [14]. Imported, unmetabolized formic acid functions as a cytoplasmic signal by binding to HilD, the master transcriptional regulator of *Salmonella* invasion, thereby preventing inhibitory fatty acids from binding and extending the activity of HilD, which leads to the derepression of invasion genes [15]. Moreover, formate utilization by respiratory enzymes such as formate dehydrogenase-N (FdnGHI) and formate dehydrogenase-O (FdoGHI) plays a key role in *Salmonella* colonization of the gut [16]. *S.* Typhimurium can source formate from both its own pyruvate-formate lyase (PflB) and from the intestinal environment [17, 18]. However, the role of endogenously produced formate in regulating *Salmonella* virulence and pathogenesis have seldom been reported [19].

In this study, we examined the role of endogenously produced formate in the pathogenic lifestyle of *S.* Typhimurium. To the best of our knowledge, we are the first to identify the role of the *pflB* gene in regulating cytosolic pH of STM, which is instrumental in maintaining the flagellar operon. Quite interestingly, in the wild-type strain with an intact formate pool, extracellular formate functions as a signalling molecule via the BarA/SirA two-component system, highlighting the distinct roles of intracellular and extracellular formate pools in regulating *Salmonella* virulence. We demonstrate that regulating the *pflB* gene in various regions of the intestine may be a strategy utilized by *Salmonella* to optimize its invasion of the distal ileum, its primary target site.

## Materials and Methods

### Bacterial strains and plasmids

The wild-type *Salmonella enterica* subspecies *enterica* serovar Typhimurium strain 14028S used in this study was generously provided by Professor Michael Hensel from the Max von Pettenkofer Institute for Hygiene and Medical Microbiology, Germany. Knockout and complement strains were generated specifically for this study. Bacterial strains were stored as glycerol stocks at -80°C and revived on fresh Luria-Bertani (LB) agar plates (HiMedia) with appropriate antibiotics as needed (50 µg/ml Kanamycin, 50 µg/ml Ampicillin, 25 µg/ml Chloramphenicol). A single colony was inoculated into a fresh LB tube with antibiotics when required and incubated at 37°C with shaking (170 rpm) to obtain an overnight culture. Strains containing the pKD46 plasmid were incubated overnight at 30°C with shaking (170 rpm). A complete list of bacterial strains and plasmids used in this study is provided in Table 1.

**Table 1:**
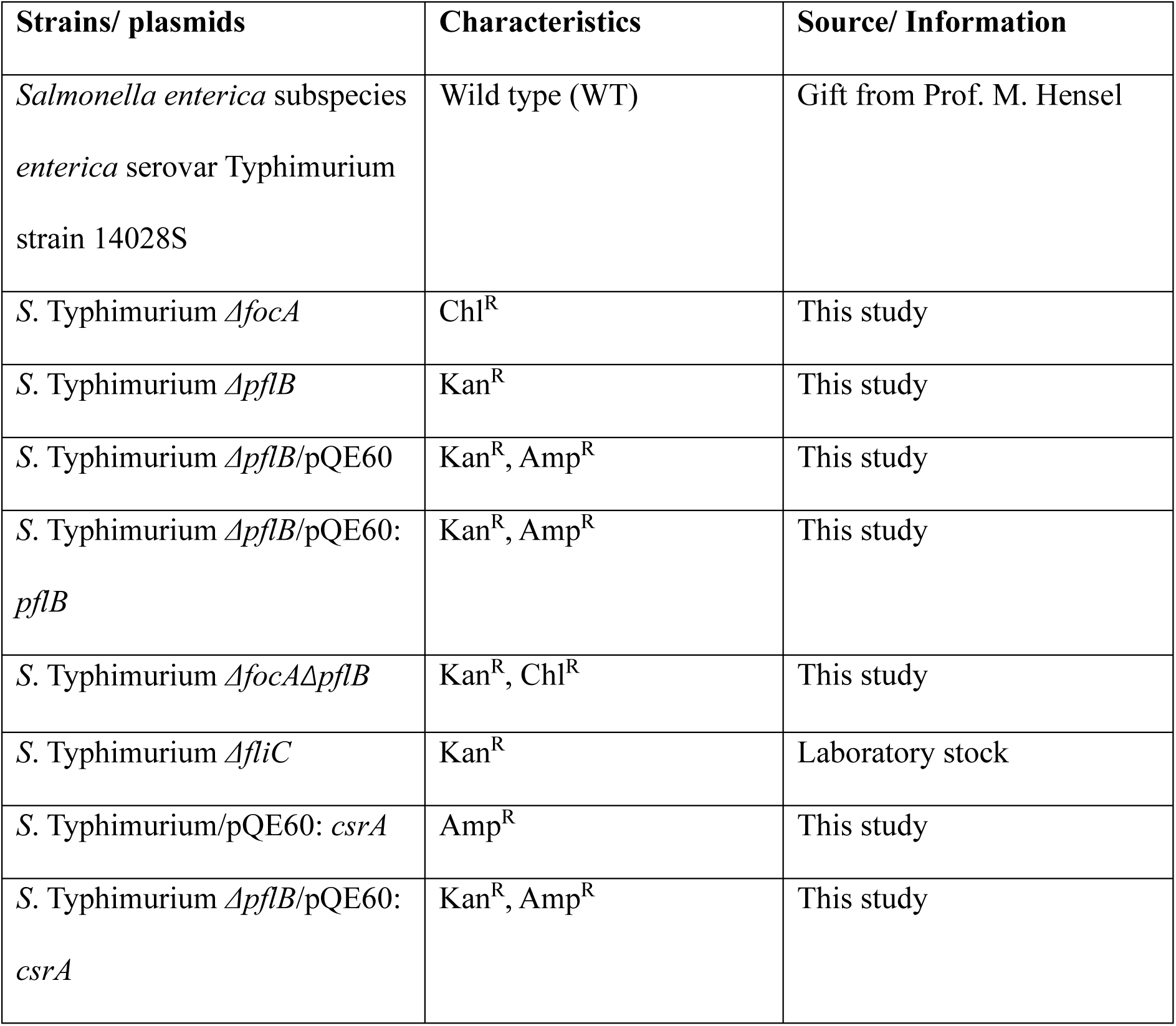
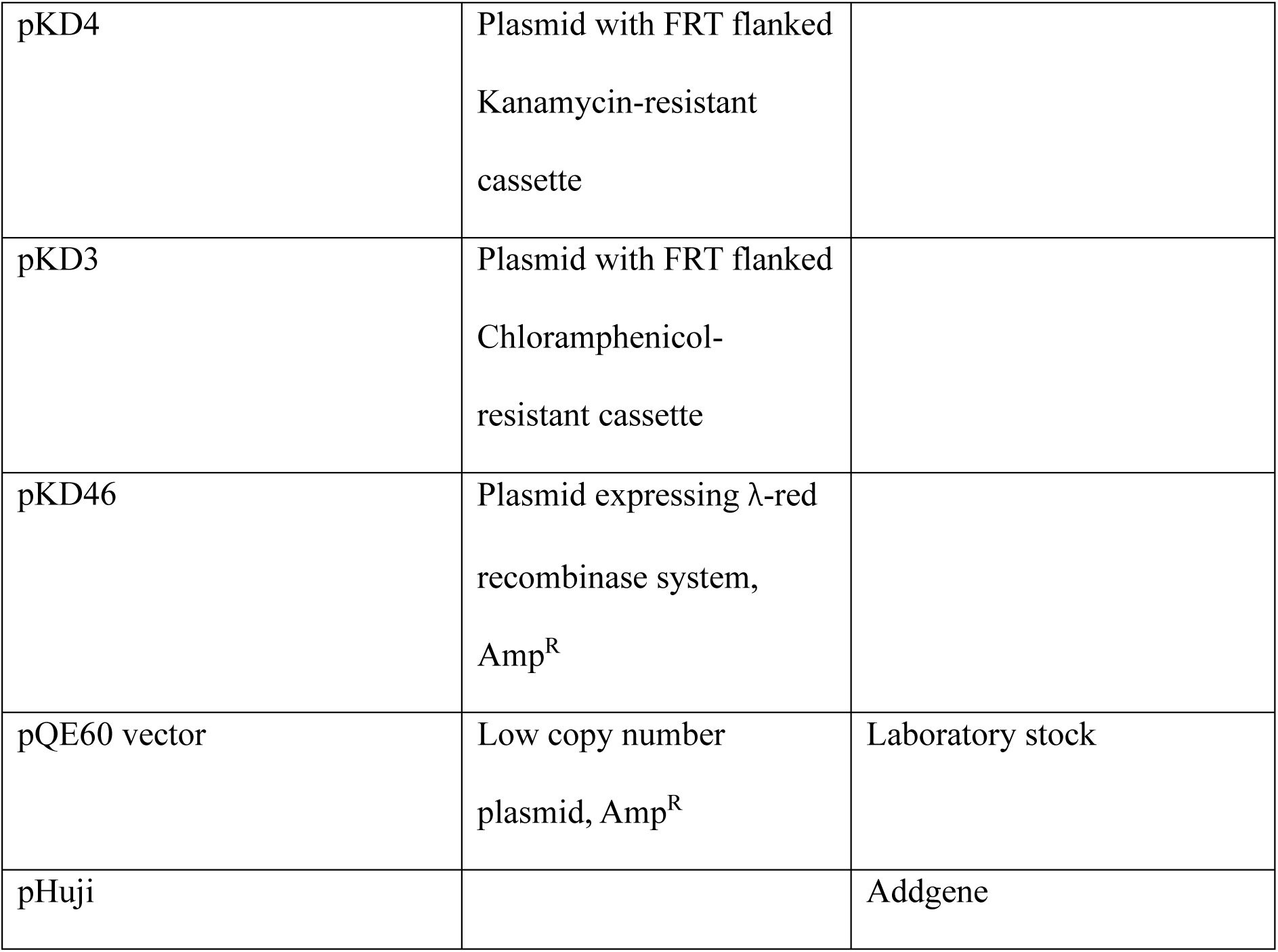
List of strains.

| Strains/ plasmids | Characteristics | Source/ Information |
| --- | --- | --- |
| *Salmonella enterica* subspecies<br>*enterica* serovar Typhimurium<br>strain 14028S | Wild type (WT) | Gift from Prof. M. Hensel |
| *S. Typhimurium* *ΔfocA* | Chl<sup>R</sup> | This study |
| *S. Typhimurium* *ΔpflB* | Kan<sup>R</sup> | This study |
| *S. Typhimurium* *ΔpflB*/pQE60 | Kan<sup>R</sup>, Amp<sup>R</sup> | This study |
| *S. Typhimurium* *ΔpflB*/pQE60:<br>*pflB* | Kan<sup>R</sup>, Amp<sup>R</sup> | This study |
| *S. Typhimurium* *ΔfocAΔpflB* | Kan<sup>R</sup>, Chl<sup>R</sup> | This study |
| *S. Typhimurium* *ΔfliC* | Kan<sup>R</sup> | Laboratory stock |
| *S. Typhimurium*/pQE60: *csrA* | Amp<sup>R</sup> | This study |
| *S. Typhimurium* *ΔpflB*/pQE60:<br>*csrA* | Kan<sup>R</sup>, Amp<sup>R</sup> | This study |
| pKD4 | Plasmid with FRT flanked<br>Kanamycin-resistant<br>cassette |  |
| pKD3 | Plasmid with FRT flanked<br>Chloramphenicol-<br>resistant cassette |  |
| pKD46 | Plasmid expressing λ-red<br>recombinase system,<br>Amp<sup>R</sup> | |
| pQE60 vector | Low copy number<br>plasmid, Amp<sup>R</sup> | Laboratory stock |
| pHuji |  | Addgene |

### Construction of the knockout strains of *Salmonella*

The knockout strains of *Salmonella* were generated using the one-step chromosomal gene inactivation technique developed by Datsenko and Wanner [20]. In brief, STM (WT) was transformed with the pKD46 plasmid, which expresses the λ-Red recombinase system under the control of an arabinose-inducible promoter. Transformed cells were selected on LB plates containing 50 µg/ml Ampicillin. To generate the knockout strains, a single colony of STM (WT)/pKD46 was inoculated into fresh LB medium with 50 µg/ml Ampicillin and 50 mM arabinose, then incubated overnight at 30°C with shaking (170 rpm). The overnight culture was then subcultured into fresh LB medium and incubated under the same conditions for 2.5 hours to reach an OD_600_ of 0.35-0.4. Electrocompetent STM (WT)/pKD46 cells were prepared by washing the bacterial pellet with chilled, autoclaved MilliQ water and 10% glycerol. Kanamycin (KanR, 1.5 kb) and Chloramphenicol (ChlR, 1.1 kb) resistance cassettes were amplified from pKD4 and pKD3 plasmids, respectively, using knockout-specific primers. The amplified resistance cassettes were then electroporated into STM (WT)/pKD46, and transformed cells were selected on LB agar plates with the appropriate antibiotics (Kanamycin or Chloramphenicol). Knockout colonies were confirmed using gene-specific intragenic primers, knockout confirmatory primers, and antibiotic-cassette-specific primers.

For generating double-knockout strains, the one-step chromosomal gene inactivation method was slightly modified. Briefly, STM Δ*pflB* was transformed with the pKD46 plasmid. An overnight culture of the transformed strain was subcultured into fresh LB medium supplemented with 50 mM arabinose and 50 µg/ml Ampicillin, then incubated at 30°C with shaking (170 rpm) for 2.5 hours until it reached an OD600 of 0.3-0.4. The chloramphenicol resistance cassette (1.1 kb), flanked by regions homologous to *focA*, was electroporated into electrocompetent STM Δ*pflB*/pKD46 cells to generate STM Δ*focAΔpflB*. The double-knockout strains were selected on LB agar plates containing Kanamycin (50 µg/ml) and Chloramphenicol (25 µg/ml). These double-knockout colonies were further confirmed with knockout confirmatory and gene-specific intragenic primers. The primers used for generating knockouts are listed in Table 2.

**Table 2:**
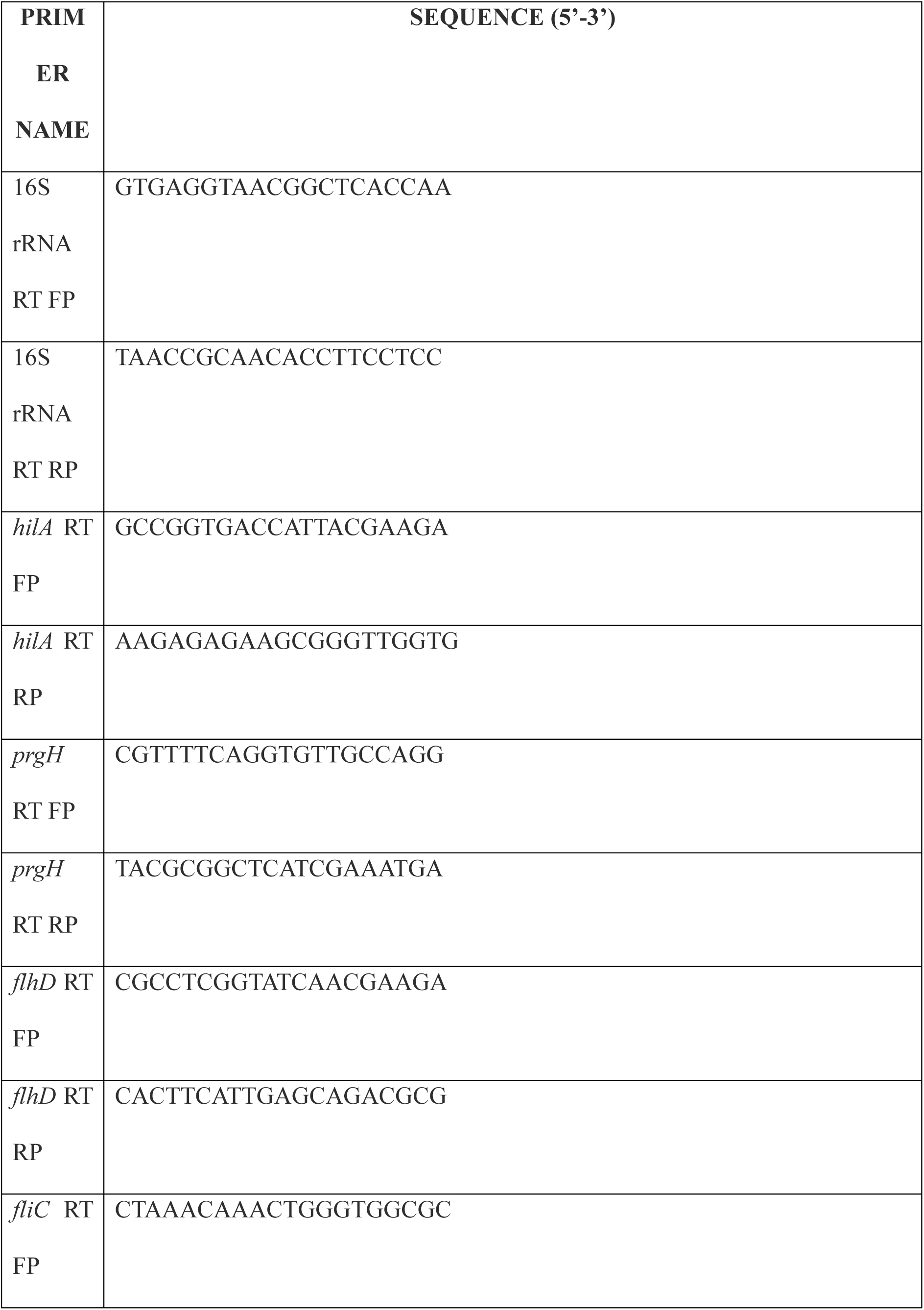
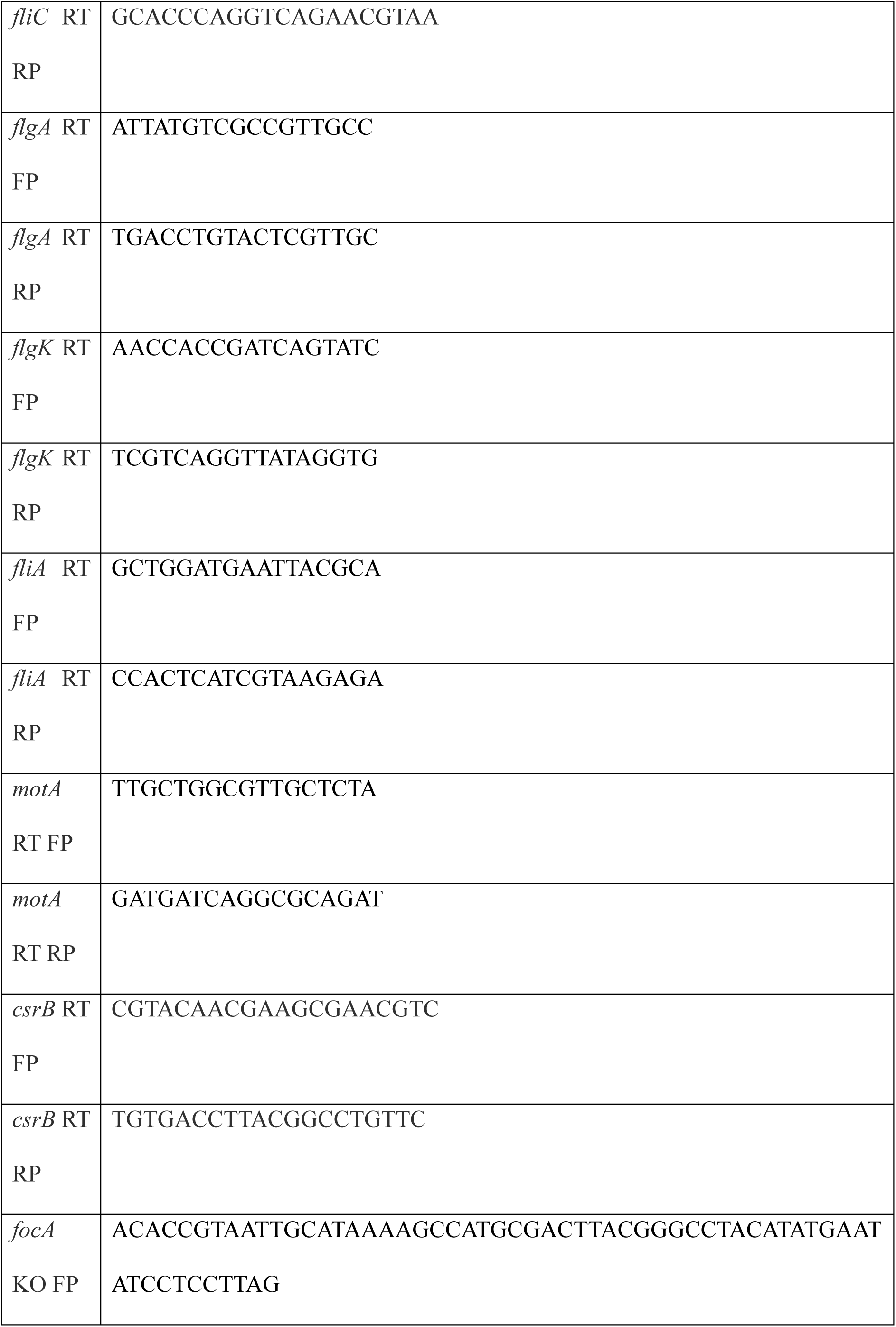
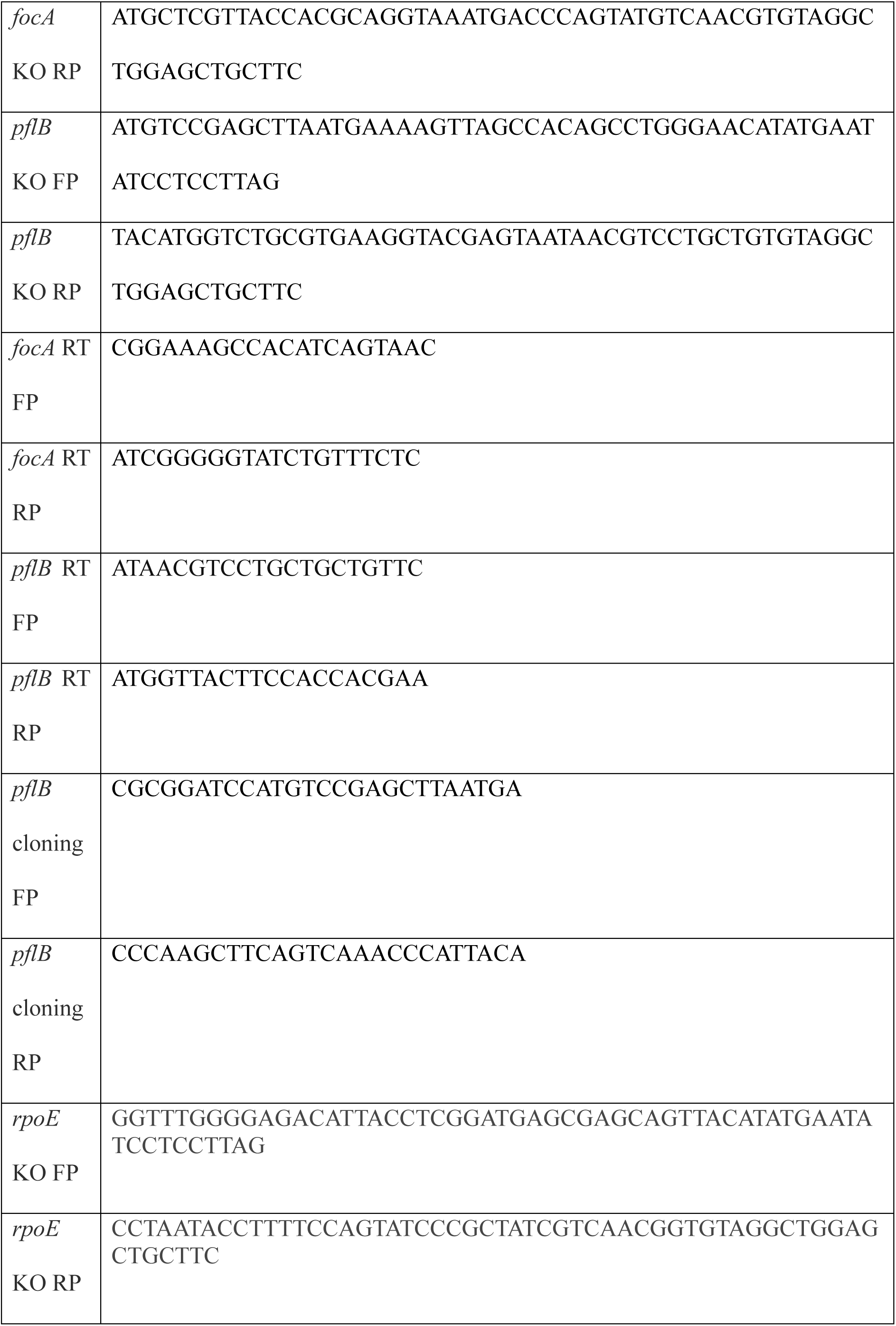
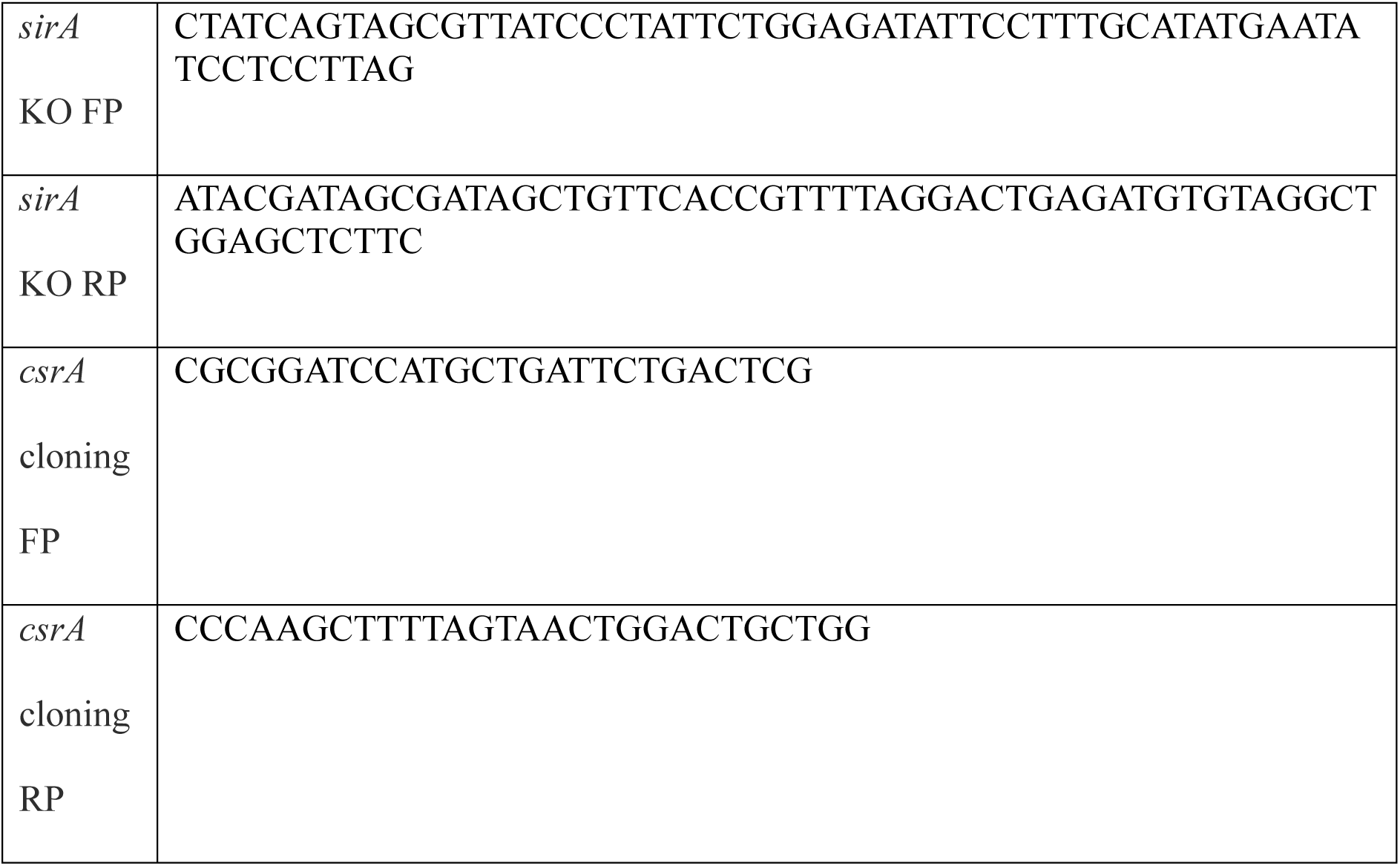
List of primers:

### Construction of complemented and overexpression strains of *Salmonella*

The *pflB* and *csrA* genes were amplified from the genomic DNA of wild-type *Salmonella* using gene-specific cloning primers. Both the colony PCR products and the empty pQE60 cloning vector were digested with BamHI-HF and HindIII-HF (NEB) at 37°C for 1 hour. After double digestion, the insert and vector were purified and ligated using T4 DNA Ligase (NEB) at 16°C overnight. The ligation product was then transformed into *Escherichia coli* TG1 using the heat shock method. Successful cloning was confirmed by double digestion of the recombinant plasmid. The confirmed recombinant plasmid was subsequently transformed into *Salmonella* strains to create complemented and overexpression strains. Primers used for generating complement strains are listed in Table 2.

### RNA Isolation and RT-qPCR

Logarithmic-phase cultures of bacterial cells was centrifuged at 6,000 rpm at 4°C, and the resulting pellet was resuspended in TRIzol reagent. The suspension was stored at -80°C until further use. RNA was isolated from the lysed supernatants using the chloroform-isopropanol method. The RNA pellet was washed with 70% ethanol and resuspended in 30 µl of DEPC-treated MilliQ water. RNA concentration was measured with a NanoDrop spectrophotometer (Thermo Fisher Scientific), and quality was assessed on a 2% agarose gel. Following this, 2 µg of RNA was treated with DNase using DNase TURBO (Thermo Fisher Scientific) at 37°C for 30 minutes. DNase was inactivated by adding 5 mM Na₂EDTA (Thermo Fisher Scientific) and incubating at 65°C for 10 minutes. cDNA synthesis was carried out using the PrimeScript RT reagent kit (Takara, CAT RR037A) according to the manufacturer’s instructions. Quantitative real-time PCR (qRT-PCR) was performed with a TB Green RT-qPCR kit (Takara) on the QuantStudio 5 Real-Time PCR system. Intragenic primers were used to examine gene expression, and expression levels were normalized to 16S rRNA. Primer sequences are listed in Table 2.

### Eukaryotic cell lines and growth conditions

The HeLa, Caco-2, HepG2, and RAW 264.7 cell lines were cultured in Dulbecco’s Modified Eagle’s Medium (DMEM, Sigma-Aldrich) supplemented with 10% Fetal Calf Serum (FCS, Gibco), at 37°C with 5% CO₂.

### Gentamicin Protection Assay and Intracellular Cell Survival Assay

Caco-2 and RAW 264.7 cells were seeded into tissue culture-treated 24-well plates. For infection of the epithelial Caco-2 cells, overnight cultures of STM WT, STM Δ*pflB*, STM Δ*pflB*/pQE60:*pflB*, and STM Δ*focAΔpflB* were subcultured in fresh LB medium, with or without supplementation of 40 mM sodium formate (Sigma). For infection of the macrophage cell line RAW 264.7, the overnight cultures were used directly. Both cell types were infected at a multiplicity of infection (MOI) of 10. The plates were centrifuged at 1000 rpm for 10 minutes, after which the cells were incubated at 37°C with 5% CO₂ for 25 minutes. Following incubation, cells were washed with 1X phosphate-buffered saline (PBS) to remove extracellular, non-adherent, and loosely adherent bacteria, and fresh media containing 100 µg/ml gentamicin was added. After a 1-hour incubation, cells were washed again with 1X PBS and exposed to 25 µg/ml gentamicin until lysis. Cell lysis was performed at 2 hours and 16 hours post-infection using 0.1% Triton X-100, and the lysates were plated onto *Salmonella*-*Shigella* (SS) agar alongside the pre-inoculum. The percent invasion and fold proliferation were determined using the following formulas:

**Percent invasion/phagocytosis= [C.F.U at 2h/ C.F.U of Pre-Inoculum] *100**

**Fold proliferation= [C.F.U at 16h/ C.F.U of 2h]**

### Adhesion assay

The adhesion assay was performed by modifying the previously reported protocol [21]. Overnight cultures (12 hours old) of STM WT, STM *ΔpflB*, and STM *ΔfliC* were subcultured into fresh LB medium, with or without supplementation of 40 mM sodium formate (Sigma). The logarithmic phase of these bacterial strains was used to infect the cells at a multiplicity of infection (MOI) of 10. The plates were centrifuged at 1000 rpm for 10 minutes, followed by incubation at 37°C with 5% CO₂ for 10 minutes. To remove extracellular, unadhered, and loosely adhered bacteria, the cells were washed twice with 1X phosphate-buffered saline (PBS). Subsequently, the cells were lysed with 0.1% Triton X-100, and the lysates were plated on Salmonella-Shigella (SS) agar alongside the pre-inoculum. The percent adhesion was determined using the following formula:

**Percent adhesion= [C.F.U at 10 mins Post Infection/ C.F.U of Pre-Inoculum] *100**

### TEM

Transmission Electron Microscopy was performed by modifying the protocol by Marathe *et al* [22]. In brief, overnight cultures of STM WT, STM *ΔpflB*, STM *ΔfocA*, STM *ΔfocAΔpflB*, STM WT/*csrA*, STM *ΔpflB*/*csrA*, STM *ΔrpoE*, and STM *ΔpflBΔrpoE* were subcultured (1:100) in fresh LB medium, with or without supplementation (+/-F), and incubated at 37°C for 2.5–3 hours under shaking conditions. The bacterial samples were then washed twice with 1X phosphate-buffered saline (PBS) and resuspended in 50 µl of 1X PBS. A 5 µl aliquot of the suspension was applied to a glow-discharged copper grid and allowed to air-dry. The bacteria were subsequently negatively stained using 1% uranyl acetate. After air-drying, the samples were imaged using Transmission Electron Microscopy (JEM-100 CX II; JEOL).

### Confocal Microscopy

Confocal Microscopy was performed by modifying the protocol by Marathe *et al* [22]. In summary, 12-hour-old stationary-phase cultures of STM WT, STM *ΔpflB*, STM *ΔfocAΔpflB*, STM WT/*csrA*, STM *ΔpflB*/*csrA*, STM *ΔrpoE*, and STM *ΔpflBΔrpoE*, with or without supplementation (+/-F), were subcultured (1:100) into fresh LB medium and incubated at 37°C for 2.5–3 hours under shaking conditions until reaching the logarithmic phase. A 100 µl aliquot of the log-phase culture was spotted onto sterile coverslips and air-dried overnight. Once completely dried, the samples were fixed with 3.5% paraformaldehyde (PFA). The PFA was washed off with 1X PBS, and the samples were stained with a rabbit-raised anti-Fli primary antibody (1:100 dilution in 2.5% BSA solution, Difco) for 3 hours at room temperature. After washing off the primary antibody with 1X PBS, the samples were incubated with Alexa-488-conjugated anti-rabbit secondary antibody (Sigma) for 45 minutes at room temperature. Before mounting the coverslips onto clean glass slides, the samples were stained with DAPI (4’,6-diamidino-2-phenylindole) at 100 µg/ml for 5 minutes. The coverslips were sealed with transparent nail polish. All images were captured using a Zeiss 880 microscope and analyzed with Zeiss ZEN Black 2012 software.

### Gas Chromatography-Mass Spectrometry

GC-MS was performed by modifying the protocol by Hughes *et al* [23]. Briefly, overnight cultures of STM WT, STM *ΔpflB*, STM *ΔfocA*, STM *ΔfocAΔpflB* were subcultured (1:100) into 100 ml of LB Broth. After 2.5-3h of incubation at 37°C under shaking conditions, 50 ml of the individual cultures were centrifuged at 6,000 rpm for 10 mins. The pellet was washed twice with 1X PBS to remove the remnants of the media. Finally, the pellet was resuspended in 1 ml of 1X PBS, and the suspension was sonicated to lyse the bacterial cells. The cell debris was removed by centrifugation. Before sonication, the CFU in the pellet had been recorded by plating.

Sodium formate (Sigma) was used as an external standard. External standards and biological samples were derivatized before the Mass Spectrometry. 2 M hydrochloric acid was added in a 1:1 ratio to the samples to protonate formate. Liquid-liquid extraction was done which extracted formic acid into ethyl acetate (Sigma-Aldrich). To remove any remaining water, the ethyl acetate extract was passed through a column of anhydrous sodium sulfate (Sigma-Aldrich). The ethyl acetate extract of formic acid was then incubated at 80 °C for 1 h with the derivatization reagent N, O-Bis(trimethylsilyl)trifluoro acetamide (Sigma Aldrich) with 1% Chlorotrimethylsilane (Fluka). The derivatized samples were then transferred to autosampler vials for analysis by Gas Chromatography Mass Spectrometry (Agilent technologies, 7890A GC, 5975C MS). The injection temperature was set to 200 °C and the injection split ratio was 1:40 with an injection volume of 1 µL. The oven temperature was set to start at 40 °C for 4 min, with an increase to 120 °C at 5 °C per min and to 200 °C at 25 °C per min with a final hold at this temperature for 3 min. Helium carrier gas had a flow rate of 1.2mL/min with constant flow mode. A 30 m × 0.25 mm × 0.40 µm DB-624 column from Agilent was used. The interface temperature was set to 220 °C. The electron impact ion source temperature was at 230 °C, with an ionization energy of 70 eV and 1235 EM volts. For qualitative experiments, selected ion monitoring (single quadrupole mode) of 103 m/z scan had been performed. The retention time for formate and deuterated formate was 5.9 min. The target and reference (qualifier) ions for formate were m/z = 103 and m/z = 75, respectively. The formate concentration of each sample was interpolated from a standard curve with the known standards. It was normalized with the CFU values of each sample to obtain the intracellular formate concentration/CFU.

To determine the formate transportation into the cell, we subcultured overnight cultures of STM WT, STM *ΔpflB*, STM *ΔfocA*, and STM *ΔfocAΔpflB* into fresh LB tubes. During the subculture, Sodium Formate was supplemented into each tube to maintain a concentration of 1M. Post 2.5-3h of incubation at 37°C under shaking conditions, the bacterial cells were removed from the media by centrifuging the culture twice at 6,000 rpm for 10 mins. A 2 µm filter was used to remove any other bacterial cells left in the spent media. Following this, a similar protocol as mentioned above was used to quantify the residual formate in the spent media. It was compared to the media where bacterial cells had not been subcultured.

### Bisbenzimide Assay

The bisbenzimide assay was performed by modifying the protocol by Roy Chowdhury *et al* [24]. Briefly, overnight cultures of STM WT, STM *ΔpflB*, and STM *ΔfliC* were subcultured into fresh LB tubes(+/-F). Logarithmic phase cultures of each condition were taken and the OD_600_ was adjusted to 0.1. To 180 µl of this bacterial suspension, 20 µl of Bisbenzimide dye (10µg/ml) was added. It was incubated at 37°C for 10 mins. The intracellular DNA-bound bisbenzimide was quantified in a TECAN 96 well microplate reader using a 346 nm excitation and 460 nm emission filter.

### Staining with DiBAC4

The DiBAC_4_ staining was performed by modifying the protocol by Roy Chowdhury *et al* [25]. Membrane potential or membrane porosity was quantified by using a membrane potential sensitive fluorescent dye bis-(1,3-dibutyl barbituric acid)-trimethylene oxonol (Invitrogen). For this purpose, overnight culture of STM WT, STM *ΔpflB*, and STM *ΔfliC* were subcultured with or without the supplementation of 40 mM Sodium Formate (Sigma). Around 4.5*10^7^ cells of the logarithmic phase culture were incubated with 1 µg/ml of DiBAC_4_ dye at 37°C for 30 mins. The percentage of DiBAC_4_-positive cells and the Median Fluorescence Intensity (MFI) of DiBAC_4_ was determined by flow cytometry (BD FACSVerse by BD Biosciences-US).

### BCECF-AM Staining

The BCECF-AM staining was performed by modifying the protocol by Roy Chowdhury *et al* [24]. The acidification of the cytosol with the change in pH of extracellular conditions was determined using a cell-permeable dual excitation ratiometric dye 2’,7’-Bis-(2-Carboxyethyl)-5-(and-6)-Carboxyfluorescein, Acetoxymethyl Ester (BCECF-AM). Briefly, the logarithmic phase cultures of STM WT, STM *ΔpflB* (+/-F) were resuspended in buffers of different pH (5,5.5,6,7). They were incubated for 2 hours at 37°C before the addition of 1 mg/ml BCECF-AM to make a final concentration of 20 µM. The cells were incubated with the dye for 30 minutes. Post-incubation, the cells were analysed by flow cytometry (BD FACSVerse by BD Biosciences, US) with 488 nm and 405 nm excitation and 535 nm excitation channels. The MFI of the bacterial cells at 488 nm and 405 nm was obtained from BD FACSuite software and the ratio of 488/405 was utilized to determine the cytosolic acidification in response to changing pH conditions.

### Intracellular pH measurement by pHuji plasmid

STM WT and STM *ΔpflB* were transformed with the plasmid pBAD:pHuji (Plasmid #61555) that encodes a pH-sensitive red fluorescent protein pHuji [26, 27]. The logarithmic cultures of STM WT/pBAD:pHuji and STM *ΔpflB*/pBAD:pHuji (+/-F) were incubated in phosphate buffers of pH 5,6,7,8,9 in the presence of 40 µM Sodium Benzoate to equilibrate the intracellular pH with the extracellular one. The resultant fluorescent intensity was plotted as a function of pH and was fitted to obtain a standard curve. The fluorescence values obtained from LB-grown cultures of the bacterial cells (+/- F) were interpolated in the standard curve to obtain the intracellular pH of the bacterial cells *in-vitro*.

### Ethics Statement for Animal Experiments

All the animal experiments have been approved by the Institutional Animal Ethics Clearance Committee (IAEC), Indian Institute of Science, Bangalore. The registration number is 48/1999/CPCSEA. The guidelines formulated by the Committee for the Purpose of Control and Supervision of Experiments on Animals (CPCSEA) were strictly followed. The ethical clearance number for this study is CAF/Ethics/853/2021.

### Mice Survival Assay

6-8 weeks old male C57BL/6 mice and Balb/c mice were acquired from the specific-pathogen free conditions of Central Animal Facility, IISc. To draw a comparison between the mortality rate of WT and mutant strains, the mice (n=10) were orally gavaged with 10^8^ cells from the stationary phase culture of each. After infection, mice were monitored every day for survival, and the data was represented as percent survival.

### Determination of bacterial burden in different organs

To understand the organ burden of the WT and mutant strain post-infection in mice, 10^7^ cells from the overnight culture of the bacterial strains were orally gavaged into 6-8 weeks old C57BL/6 mice. 5 days post-infection, the mice were ethically euthanized, and the liver, spleen, and brain were acquired. The organs were homogenized, and appropriate dilutions of the homogenate were plated on *Salmonella-Shigella* (SS Agar). The data was represented was log_10_CFU/gm-wt.

### Invasion Assay in mice

Invasion assay in mice was performed by modifying the protocol by Chandra *et al* [28]. Briefly, 10^7^ cells from the overnight culture of the bacterial cells were orally gavaged into 6-8 weeks old male C57BL/6 mice. 6 hours post-infection, the mice were ethically euthanized, and the intestine was acquired. It was homogenized, and appropriate dilutions of the homogenate were plated on *Salmonella-Shigella* (SS Agar). Corresponding fecal matter was also collected, and it was also plated post homogenization. The data was represented was log_10_CFU/gm-wt.

### Intraperitoneal infection of mice

Intraperitoneal infection of mice was performed with 10^3^ CFU from the overnight cultures of WT and mutant strains. 2 days post-infection, the mice were ethically euthanized, and the liver and the spleen were obtained. The organ burden of the bacterial strains was determined by the same protocol as above.

### Statistical Analysis

Each experiment has been repeated 1 to 8 times independently as mentioned in the figure legends. The statistical analyses were performed by a unpaired Student’s t-test or two-way ANOVA multiple comparison test as denoted in the figure legends. The data obtained from the mice experiments were analyzed using the Mann-Whitney *U* test. The *p*-values below 0.05 were considered significant. The results are indicated as Mean ± SD or Mean ± SEM as mentioned in the figure legends. All the data were plotted and analysed using GraphPad Prism 8.4.2.

## Results

### The deletion of the *pflB* gene led to decreased invasion efficiency in the Caco-2 cell line, which corresponded to a reduced organ burden in C57BL/6 mice five days after infection through oral gavaging

To investigate the role of endogenous formate levels in *Salmonella* virulence and pathogenesis, we created knockout strains of *pflB* and *focA* using the one-step gene inactivation technique outlined by Datsenko and Wanner [20]. FocA is the formate transporter in *Salmonella* Typhimurium that functions as a passive export channel at high external pH and switches to an active importer of formate at low pH [29]. The deletion of either gene did not impact the *in vitro* growth of *Salmonella* in either complex LB media or M9 Minimal Media (**Figure S1 A-B**). Since the *focA* and *pflB* genes are part of the same operon, we investigated whether deleting one gene affected the expression of the other [30]. Our study revealed that deleting the upstream *focA* gene resulted in decreased expression of the downstream *pflB* gene (**Figure S1C**). However, deletion of *pflB* increased the expression of *focA* (**Figure S1D**).

To confirm that deleting *pflB* and *focA* indeed depleted the native formate pool of the cell, we collected lysates from logarithmic phase cultures of all strains and quantified the formate concentration using Gas Chromatography-Mass Spectrometry (GC-MS). After normalizing by CFU, we observed that there was a significant reduction in the relative intracellular formate concentrations upon deletion of *pflB*, whereas deletion of *focA* did not cause a notable change. There was a significantly reduced intracellular formate level in STM *ΔfocAΔpflB* (**Figure 1A**). This finding suggested the possibility of an alternative formate transporter. To investigate further, we incubated secondary cultures of all strains in formate-supplemented media and measured formate concentration in the spent media after 3 hours by GC-MS. We found a significant reduction in formate concentration in the spent media of STM *ΔfocA* compared to the control, indicating the presence of an alternative formate importer (**Figure 1B**). Since STM Δ*focA* retained intracellular formate levels similar to STM WT and our data suggested the presence of an alternate formate transporter, we focused our study on STM Δ*pflB*, where we could successfully deplete the intracellular formate pool.

**Figure 1:**
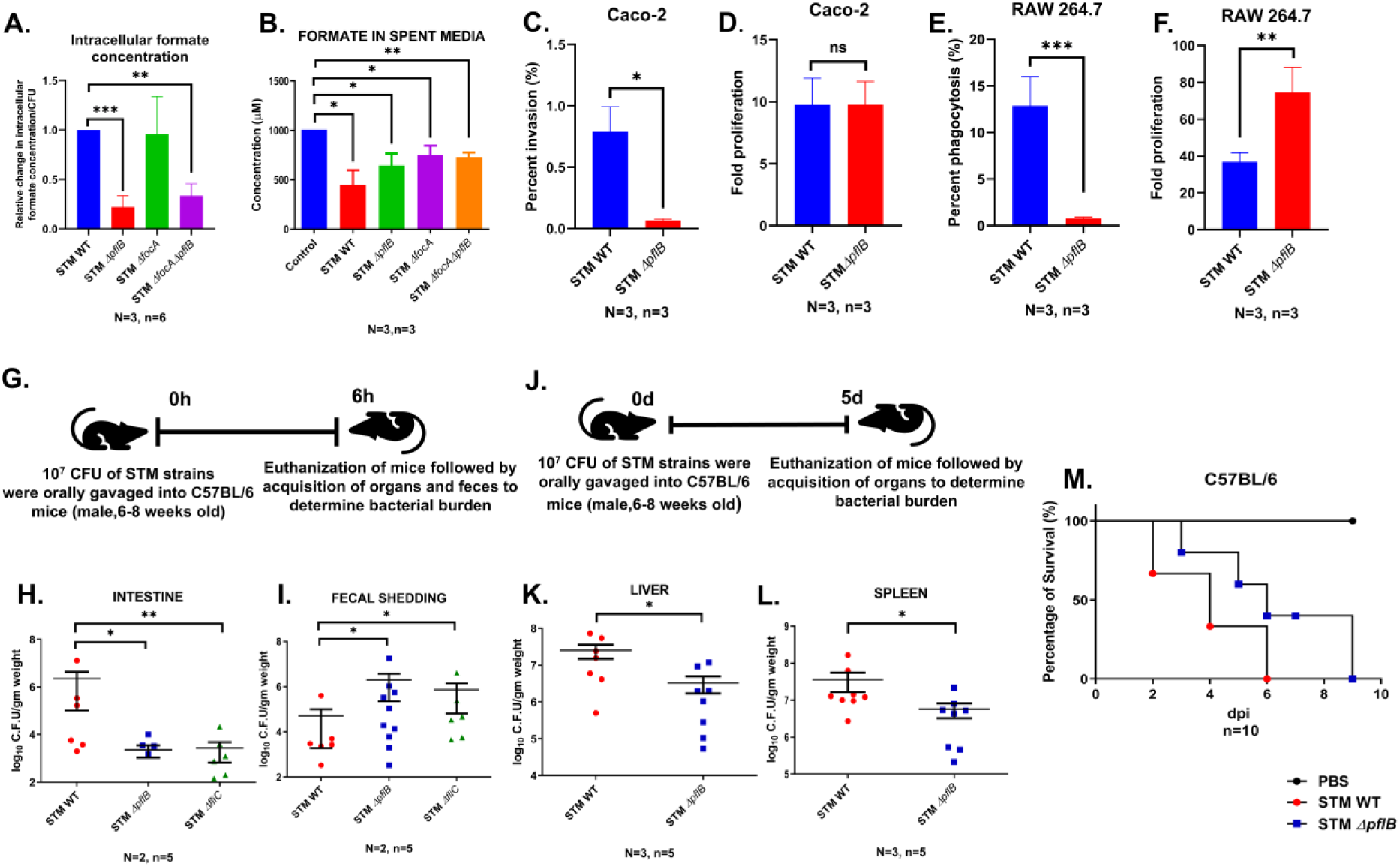
The deletion of the *pflB* gene led to decreased invasion efficiency in the Caco-2 cell line, which corresponded to a reduced organ burden in C57BL/6 mice five days after infection through oral gavaging. A. GC-MS mediated quantification of relative intracellular formate in STM WT, STM *ΔpflB*, STM *ΔfocA*, STM *ΔfocAΔpflB*. Data is represented as Mean+/-SEM of N=3, n=6. B. GC-MS mediated quantification of residual formate in the spent media post incubation by STM WT, STM *ΔpflB*, STM *ΔfocA*, STM *ΔfocAΔpflB*. Data is represented as Mean+/-SEM of N=3, n=3. C. Percent invasion of STM WT and STM *ΔpflB* in Caco-2 cell line. Data is represented as Mean+/-SEM of N=3, n=3. D. Fold proliferation of STM WT and STM *ΔpflB* in Caco-2 cell line. Data is represented as Mean+/-SEM of N=3, n=3. E. Percent phagocytosis of STM WT and STM *ΔpflB* in RAW 264.7 cell line. Data is represented as Mean+/-SEM of N=3, n=3. F. Fold proliferation of STM WT and STM *ΔpflB* in RAW 264.7 cell line. Data is represented as Mean+/-SEM of N=3, n=3. G. Schematic showing the protocol followed for invasion assay in C57BL/6 mice H. Bacterial burden in the intestine post 6h of infecting the mice with STM WT and STM *ΔpflB* via oral gavaging. Data is represented as Mean+/-SEM of N=2, n=5. I. Bacterial burden in the feces of mice post 6h of infection with STM WT and STM *ΔpflB* via oral gavaging. Data is represented as Mean+/-SEM of N=2, n=5. J. Schematic showing the protocol followed for determining the organ burden of STM WT and STM *ΔpflB* 5 days post oral gavaging K. Organ burden of STM WT and STM *ΔpflB* in liver 5 days post oral gavaging. Data is represented as Mean+/-SEM of N=3, n=5. L. Organ burden of STM WT and STM *ΔpflB* in spleen 5 days post oral gavaging. Data is represented as Mean+/-SEM of N=3, n=5. M. Survival rates of C57BL/6 mice post infection with STM WT and STM *ΔpflB* through oral route. (Unpaired Student’s t-test for column graphs, Two-way Anova for grouped data, Mann-Whitney U-test for animal experiment data (**** p < 0.0001, *** p < 0.001, ** p<0.01, * p<0.05))

Intracellular Cell Survival Assay (ICSA) using the human colorectal carcinoma cell line Caco-2 showed that STM *ΔpflB* exhibited reduced invasion efficiency compared to STM WT, but it had similar fold proliferation to the latter (**Figure 1C**). The invasion deficiency in STM *ΔpflB* was also rescued by complementing the gene with an exogenous pQE60 plasmid (**Figure S1E**).

In the murine macrophage cell line RAW 264.7, STM *ΔpflB* showed a decreased phagocytosis and hyperproliferation compared to STM WT (**Figure 1E, F**).

To determine if the invasion deficiency observed in STM *ΔpflB* was consistent in an animal model, we orally gavaged 10^7^ CFU of STM WT and STM Δ*pflB* into 6–8-week-old male C57BL/6 mice. Adhesion-deficient STM *ΔfliC* served as a control. Six hours post-infection, we observed a reduced bacterial load in the mouse intestine for STM Δ*pflB*, which was associated with increased faecal shedding compared to STM WT (**Figure 1G, H, I**). These findings were consistent with the results for the control strain STM *ΔfliC*, which is known to have reduced adhesion and is cleared more rapidly from the host [31].

To further understand the role of *pflB* deletion in overall *Salmonella* pathogenesis in *in-vivo* mouse model, we orally gavaged 10^7^ CFU of bacteria in 6-8 weeks old male C57BL/6 mice and euthanized the mice 5 days post-infection (dpi) to determine the bacterial burden in the liver, the spleen, and the brain (**Figure 1J, K, L**). We found a significantly reduced bacterial burden in the liver and the spleen when infected with STM *ΔpflB* as compared to infection with STM WT (**Figure 1K, L**). We also found that C57BL/6 mice infected with STM *ΔpflB* had a better survival rate as compared to mice infected with STM WT (**Figure 1M**). We subjected C57BL/6 mice to an intra-peritoneal infection with 10^3^ CFU of both the WT and the knockout strains. Under these conditions, we observed no significant change in the organ burden of STM WT and STM *ΔpflB* 3dpi (**Figure S2A-D**). All these data suggested that the reduced organ burden of STM *ΔpflB* upon oral gavaging is due to its attenuated adhesion in the gastrointestinal tract, which promoted a faster clearance from the host body.

### STM *ΔpflB* was deficient in flagella and the deficiency was partially recovered upon supplementation of 40 mM sodium formate

*Salmonella* Typhimurium uses adhesins, such as flagella, to attach to host cells during infection [32-34]. Following successful adhesion, it employs the *Salmonella* Pathogenicity Island-1 (SPI-1) machinery to invade non-phagocytic epithelial cells [35-37]. Our previous findings indicated that STM Δ*pflB* has reduced invasion efficiency in Caco-2 cells. We opted to proceed by supplementing exogenous formate to STM *ΔpflB* to determine whether the effects observed from depleting intracellular formate could indeed be reversed through the addition of extracellular formate. In our further studies, we used a concentration of 40 mM (indicated as F henceforth), as it yielded maximal expression of the SPI-1 regulatory gene *hilA* in STM WT in a concentration-dependent analysis ranging from 0 to 50 mM (**Figure S3A**). Interestingly, we observed that STM Δ*pflB* exhibited increased expression of *prgH*, which encodes a structural protein of the SPI-1 apparatus (**Figure 2A**). This expression was further elevated in STM *ΔpflB*+F. Additionally, STM WT+F showed significantly higher *prgH* expression compared to STM WT, consistent with the findings of Huang *et al* [17].

**Figure 2:**
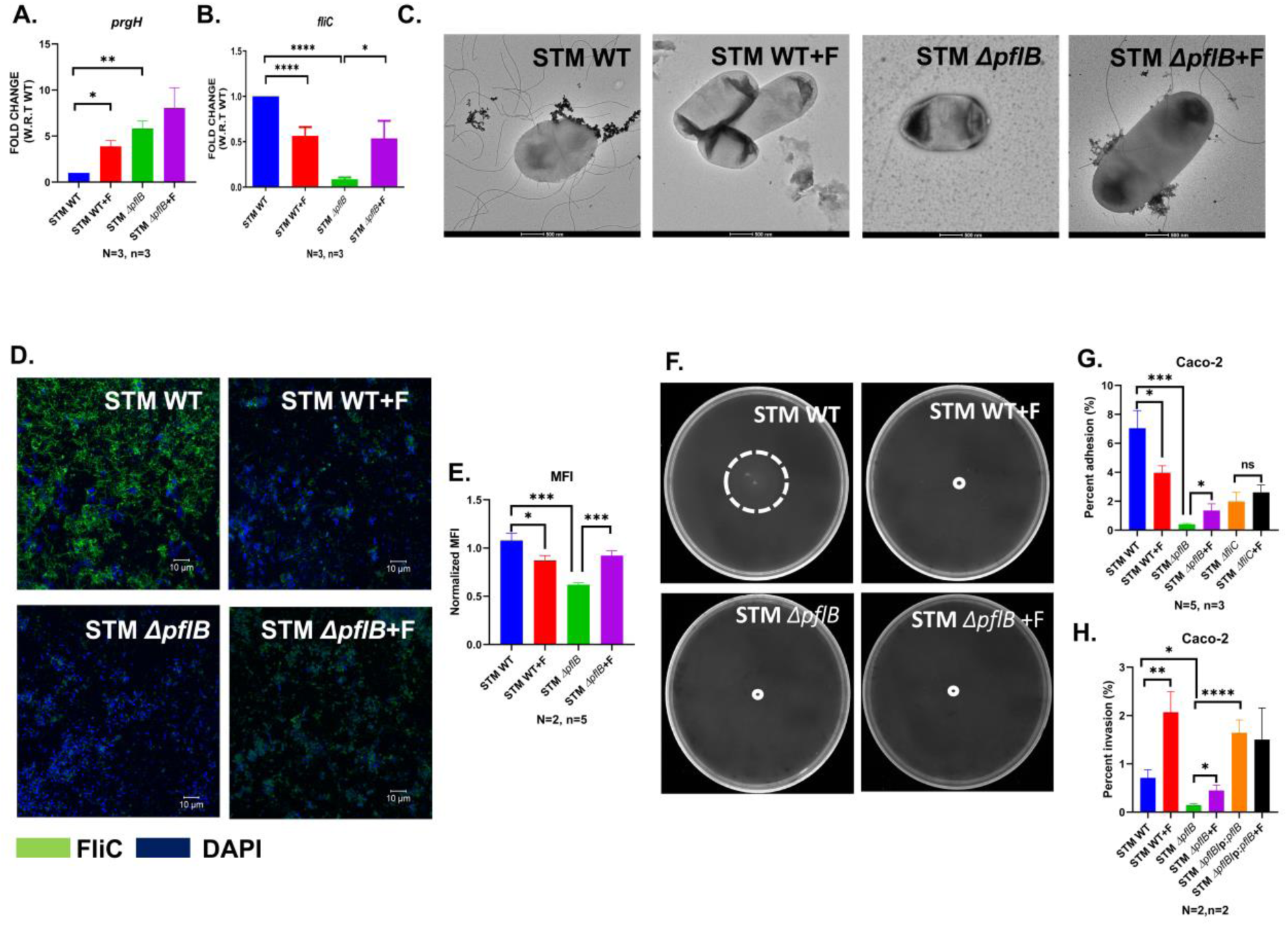
STM *ΔpflB* was deficient in flagella and the deficiency was recovered upon supplementation of formate. A. RT-qPCR mediated study of expression profile of *prgH* in the logarithmic phase cultures of STM WT, STM *ΔpflB* (+/-F). Data is represented as Mean+/-SEM of N=3, n=3. B. RT-qPCR mediated study of expression profile of *fliC* in the logarithmic phase cultures of STM WT, STM *ΔpflB* (+/-F). Data is represented as Mean+/-SEM of N=2, n=3. C. TEM assisted visualization of the flagellar structures in STM WT, STM *ΔpflB* (+/-F) D. Confocal microscopy assisted visualization of the flagellar structures in STM WT, STM *ΔpflB* (+/-F). Images are representative of N=2, n=5. E. Normalized Median Fluorescence Intensity (MFI), i.e.., MFI of Secondary antibody for Rabbit generated Anti-fli antibody/ MFI for DAPI in confocal microscopy assisted visualization of the flagellar structures in STM WT, STM *ΔpflB* (+/-F). Data is represented as Mean+/-SEM of N=2, n=5. F. Swim motility of STM WT and STM *ΔpflB* on 0.25% Agar containing LB plates G. Percent adhesion of STM WT, STM *ΔpflB*, STM *ΔfliC*(+/-F) in Caco-2 cell line. Data is represented as Mean+/-SEM of N=5, n=3. H. Percent invasion of STM WT, STM *ΔpflB*, STM *ΔpflB/*pQE60:*pflB* (+/-F) in Caco-2 cell line. Caco-2 cell line. Data is represented as Mean+/-SEM of N=2, n=2 (Unpaired Student’s t-test for column graphs, Two-way Anova for grouped data, Mann-Whitney U-test for animal experiment data (**** p < 0.0001, *** p < 0.001, ** p<0.01, * p<0.05))

We also found that STM *ΔpflB* had lower *fliC* expression, which was partially restored under +F conditions. Formate supplementation reduced *fliC* expression in STM WT as well, compared to the untreated control (**Figure 2B**). We confirmed this via Transmission Electron Microscopy (TEM) as well, where we found that STM WT+F had lesser flagella compared to untreated control. STM *ΔpflB* had a reduced flagellar number, which was partially restored upon supplementation of F (**Figure 2C**). Quantitative analysis of normalized MFI in confocal microscopy and adhesion assays in the Caco-2 cell line further validated the findings from qRT-PCR and TEM (**Figure 2 D-F**). STM WT+F also showed lesser motility on swim agar plate compared to STM WT, however, we did not see a recovery in motility of STM *ΔpflB* upon F supplementation (**Figure 2 G**). This indicates that while F supplementation in STM *ΔpflB* may restore the transcription level of *fliC* and its protein expression, it does not lead to a recovery in flagella-mediated motility. The partially restored adhesion observed in STM Δ*pflB*+F in the Caco-2 cell line could be attributed to the increased surface hydrophobicity of the flagellar filaments, resulting from the recovery in protein levels. This enhanced hydrophobicity promotes stronger interactions for effective adhesion to eukaryotic host cells [38]. F supplementation in STM *ΔpflB* was also associated with an improved invasion in Caco-2 cell line. Complementation of the *pflB* via an extragenous plasmid in STM *ΔpflB* also led to an improved percent invasion compared to the knockout strain. In line with our previous data, we also found that STM WT+F had a higher percent invasion compared to STM WT (**Figure 2H**).

Further experimentation involved incubating secondary cultures of STM WT and STM Δ*pflB* in varying concentrations of formate. Notably, formate significantly inhibited *fliC* expression in STM WT starting from a concentration of 1 mM, while a concentration as low as 0.125 mM was sufficient to restore reduced *fliC* expression in STM Δ*pflB* (**Figure S3B**). Additionally, 10 mM formate significantly upregulated *hilA*—the master regulator of SPI-1—in both strains (**Figure S3C**).

The double knockout strain STM Δ*focAΔpflB* exhibited similar growth kinetics as STM WT in LB broth and M9 Minimal Media (**Figure S4A, B**). However, due to its depleted formate pool, it was also deficient in flagella, with partial recovery in anti-Fli MFI and adhesion in Caco-2 cell line when incubated with formate (**Figure S4 C, D, E**). In line with our previous findings, we observed that oral gavaging of STM *ΔfocA* led to a similar bacterial burden in the liver and the spleen as STM WT 5 dpi. STM *ΔfocAΔpflB* showed lesser organ burden in the liver and spleen of the C57 mice model (**Figure S4 F**).

### STM *ΔpflB* had a higher intracellular pH and higher membrane depolarization, both of which were recovered upon F supplementation

Previous studies in *Escherichia coli* indicated that formate translocation through the FocA channel and its metabolism via the FHL-1 (Formate Hydrogen Lyase-1) complex are crucial for maintaining cellular pH homeostasis during anaerobic growth or fermentation [39-45]. This raised the question of whether formate synthesis during aerobic growth in *Salmonella* Typhimurium could also contribute to maintaining intracellular pH (ipH). Additionally, weak acids like acetate and benzoate can cross the cell membrane in their neutral forms and subsequently dissociate, thereby reducing ipH [46]. Based on this, we also hypothesized that the endogenous formate pool might play a role in maintaining intracellular pH (ipH). To investigate how changes in the surrounding media’s pH affect intracellular pH, we incubated STM WT and STM *ΔpflB* (+/- F) in phosphate buffers at pH levels 5.5, 6, and 7. After incubation, we stained the cells with the pH-sensitive dye BCECF-AM. A higher 488/405 nm fluorescence ratio for STM *ΔpflB* under all pH conditions indicated a higher ipH in the knockout strain compared to the WT (**Figure 3A**).

**Figure 3:**
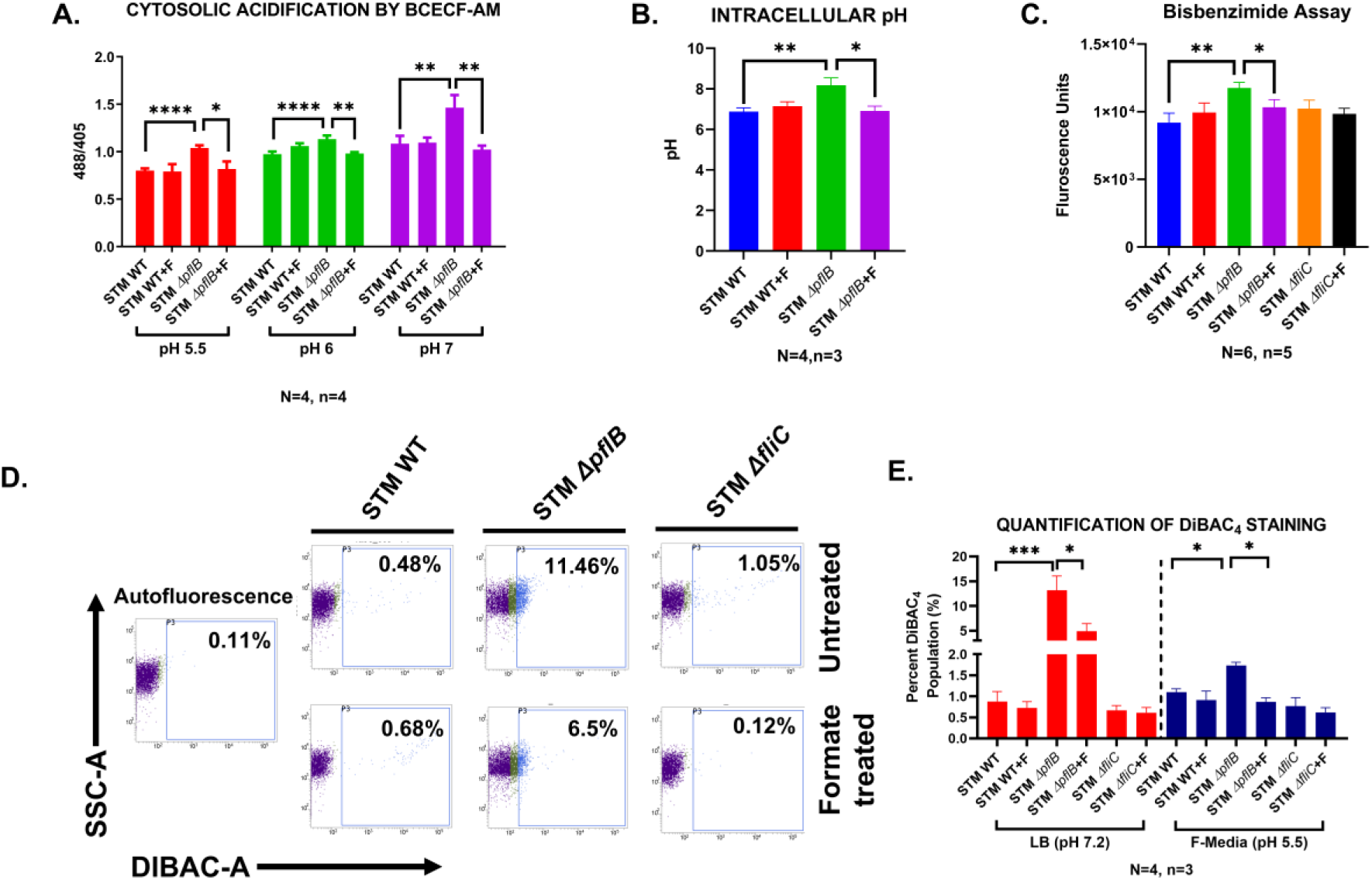
STM ***Δ****pflB* had a higher intracellular pH and higher membrane depolarization, both of which were recovered upon formate supplementation. A. BCECF-AM dye mediated elucidation of the relative ipH in the logarithmic cultures of STM WT, STM *ΔpflB* (+/-F) in phosphate buffers of pH 5.5, 6, and 7. Data is represented as Mean+/-SEM of N=4, n=4. B. Elucidation of the ipH in the logarithmic cultures of STM WT/pBAD-pHuji and STM *ΔpflB*/pBAD-pHuji (+/-F). Data is represented as Mean+/-SEM of N=4, n=3. C. Bisbenzimide assay to quantify the membrane damage in the logarithmic cultures of STM WT, STM *ΔpflB,* STM *ΔfliC* (+/-F). Data is represented as Mean+/-SEM of N=6, n=5. D. Representative FACS percent positive and histogram profiles of DiBAC_4_ staining in the logarithmic cultures of STM WT, STM *ΔpflB,* STM *ΔfliC* (+/-F). Data is representative of N=4, n=3. E. Flow cytometry obtained quantification of percent-DiBAC_4_ positive cells in the logarithmic cultures of STM WT, STM *ΔpflB,* STM *ΔfliC* (+/-F) incubated in LB media (pH 7.2) and F-media (pH 5.5). Data is represented as Mean+/-SEM of N=4, n=3. (Unpaired Student’s t-test for column graphs, Two-way Anova for grouped data, Mann-Whitney U-test for animal experiment data (**** p < 0.0001, *** p < 0.001, ** p<0.01, * p<0.05))

To further confirm these results, we transformed STM WT and STM *ΔpflB* with the plasmid pBAD-pHuji, which encodes a pH-sensitive red fluorescent protein. The higher ipH observed in STM *ΔpflB*/pBAD-pHuji confirmed that *pflB* plays a significant role in maintaining cytosolic pH in *Salmonella* Typhimurium. Consistent with the BCECF-AM staining results, formate did not affect the ipH of STM WT/pBAD-pHuji (**Figure 3B**). These findings suggest the possibility that extracellular and intracellular formate pools might independently regulate the expression of *fliC* and *hilA* through distinct mechanisms.

One of the causes of membrane depolarization in biological systems is disrupted pH homeostasis [47]. To investigate membrane potential in all strains with and without formate supplementation, we analysed the exponential cultures of STM *ΔpflB* in LB. These cultures showed a higher population of DiBAC_4_-positive cells, indicating increased membrane depolarization. Supplementation with formate successfully reduced membrane depolarization in STM *ΔpflB*, but it had no effect on membrane polarization in STM WT (**Figure 3D, E**). Similar results were observed when cells were stained with the DNA-binding fluorescent dye bisbenzimide (**Figure 3C**).

Furthermore, the extent of membrane depolarization in STM *ΔpflB* was reduced when stationary-phase bacteria were sub cultured into acidic F-media (pH 5) (**Figure 3E**). The H⁺ ions in the acidic F-media may help counteract the disrupted cytosolic acidification in the *pflB* knockout strain, thereby restoring membrane polarization.

These findings suggest that the intracellular formate pool is crucial for maintaining intracellular pH (ipH) in *Salmonella* Typhimurium, which in turn supports cell membrane polarization. We also explored whether the elevated ipH in the *pflB* knockout strain contributes to disrupted flagellar apparatus synthesis. Incubation of STM *ΔpflB* in acidic LB medium (pH 6) led to an increase in the *fliC* transcript level (**Figure S5 A**), indicating that higher ipH is responsible for these downstream effects.

### STM *ΔpflB* had a higher transcript level of *csrB*, raising the possibility of flagella inhibition via the *csr* sRNA system

To investigate how formate regulates the flagellar operon, we measured the transcript levels of several genes across different classes, including class I (*flhD*), class II (*flgA*, *fliA*), and class III (*flgK*, *motA*) (**Figure 4A**) [48-56]. We discovered that in the formate-deficient *pflB* knockout strain, inhibition of the flagellar operon begins at the level of class I genes or the genes that regulate class I genes. The transcription of the *flhDC* promoter is controlled by various factors, including OmpR, Crp, HdfR, H-NS, QseBC, FliA, LrhA, and others [50, 57-59]. One of the most extensively studied mechanisms of post-transcriptional regulation of *flhDC* involves the Csr system. The Csr system consists of the RNA-binding protein CsrA and the small regulatory RNAs (sRNAs) *csrB* and *csrC* [60]. CsrA is a 61-amino-acid protein that binds to the mRNA of target genes, thereby either inhibiting or enhancing their translation [61]. CsrA can bind to the *flhD* transcript and stabilize it by preventing RNase E cleavage at the 5′ end [62]. The activity of the CsrA protein is antagonized by small regulatory RNAs *csrB* and *csrC*. A single *csrB* RNA molecule can bind up to 18 CsrA molecules, whereas a single *csrC* molecule can bind up to 9 CsrA molecules [63-65]. In *S.* Typhimurium, CsrA has been seen to regulate motility, virulence, carbon storage, and secondary metabolism [66, 67]. The transcription of *csrB* and *csrC* is regulated by the BarA/SirA two-component systems (TCS) orthologues in *E. coli* and *S.* Typhimurium[68-71].

**Figure 4:**
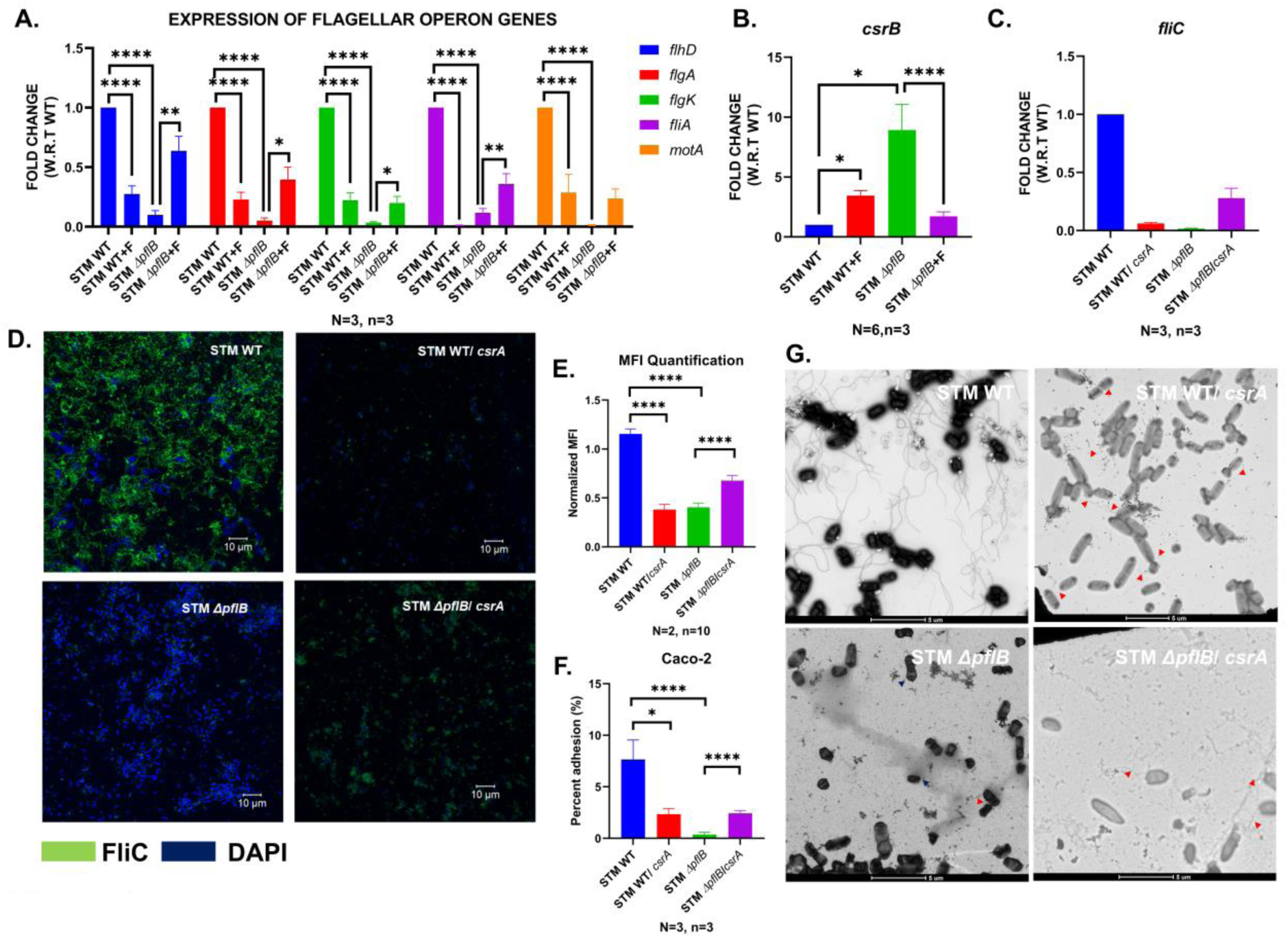
STM *ΔpflB* had a higher transcript level of *csrB*, raising the possibility of flagella inhibition via the *csr* sRNA system. A. RT-qPCR mediated expression profile of the flagellar operon genes *flhD*, *flgA*, *flgK*, *fliA*, *motA* in the logarithmic cultures of STM WT, STM *ΔpflB (+/-F).* Data is represented as Mean+/-SEM of N=3, n=3. B. RT-qPCR mediated expression profile of the sRNA *csrB* in the logarithmic cultures of STM WT, STM *ΔpflB (+/-F).* Data is represented as Mean+/-SEM of N=6, n=3. C. RT-qPCR mediated expression profile of the *fliC* in the logarithmic cultures of STM WT and STM *ΔpflB* transformed with an pQE60*: csrA* under an IPTG-inducible system. Data is represented as Mean+/-SEM of N=3, n=3. D. Confocal microscopy assisted visualization of the flagellar structures in STM WT/pQE60: *csrA*, STM *ΔpflB*/pQE60: *csrA*. Images are representative of N=2, n=10. E. Normalized Median Fluorescence Intensity (MFI), i.e.., MFI of Secondary antibody for Rabbit generated Anti-fli antibody/ MFI for DAPI in confocal microscopy assisted visualization of the flagellar structures in STM WT/pQE60: *csrA*, STM *ΔpflB*/pQE60: *csrA*. Data is represented as Mean+/-SEM of N=2, n=10 F. Percent adhesion of STM WT/pQE60: *csrA* and STM *ΔpflB*/pQE60: *csrA* in Caco-2 cell line. Data is represented as Mean+/-SEM of N=3, n=3. G. TEM assisted visualization of the flagellar structures in STM WT/pQE60: *csrA*, STM *ΔpflB*/pQE60: *csrA.* Data is representative of N=2, n=10. (Unpaired Student’s t-test for column graphs, Two-way Anova for grouped data, Mann-Whitney U-test for animal experiment data (**** p < 0.0001, *** p < 0.001, ** p<0.01, * p<0.05))

We observed an increase in *csrB* transcript levels in STM *ΔpflB*, which decreased in STM *ΔpflB*+F (**Figure 4B**). We also observed a reduction in *csrA* transcript levels in STM *ΔpflB,* which was recovered upon incubation in F supplemented LB or LB with acidic pH (pH 6) (**Figure S5B**). Since *csrB* directly binds to CsrA and inhibits its activity, we decided to clone *csrA* into the extragenous plasmid pQE60 and transform this recombinant plasmid into STM *ΔpflB*. Overexpressing *csrA* through IPTG induction partially restored the reduced expression of *fliC* in STM *ΔpflB* (**Figure 4C**). CsrA is known to bind and stabilize the *flhD* transcript. However, we speculate that an excess of CsrA molecules in the cell could potentially stabilize *flhD* to the point of inhibiting its translation. This may explain why *csrA* overexpression in STM WT led to a decrease in *fliC* expression. Overexpression of *csrA* mediated downregulation of motility has been demonstrated in *Bacillus subtilis* and *Clostridium difficile* as well [72, 73]. The increase in flagellar numbers upon *csrA* overexpression in STM *ΔpflB* was visualized and quantified using TEM and confocal microscopy (**Figure 4D**, **E, G**). This finding was corroborated by an adhesion assay with the Caco-2 cell line, where STM *ΔpflB*/p: *csrA* showed a higher adhesion percentage compared to STM *ΔpflB* (**Figure 4F**).

### The extracytoplasmic sigma factor RpoE enhances the expression of csrB, which in turn limits the expression of *fliC*

In *E. coli*, RpoE (σ^E^) functions as an extracytoplasmic stress factor essential for cell viability and regulating the response to membrane stress[74]. The membrane-bound anti-sigma factor RseA binds to σ^E^, aided by the periplasmic protein RseB [75, 76]. In the absence of envelope stress, RseA binding inhibits σ^E^ from interacting with RNA polymerase [76-79]. When envelope stress occurs, the proteolytically unstable RseA is quickly degraded, releasing free σ^E^ molecules to carry out downstream functions. These functions include activating genes involved in the biogenesis, transport, and assembly of lipopolysaccharides (LPS), phospholipids, and outer membrane proteins (OMPs), as well as proteases and chaperones that maintain or repair outer membrane (OM) integrity [76]. Recent studies have demonstrated that σ^E^ also indirectly activates the transcription of *csrB*/*C* sRNAs from σ^70^ promoters [80, 81]. These findings have been established in *E. coli*. Deletion of *rpoE* in STM showed a reduction in *csrB* transcript levels (∼0.588), suggesting that a similar mechanism may operate in STM as well **(Figure 5A)**.

**Figure 5:**
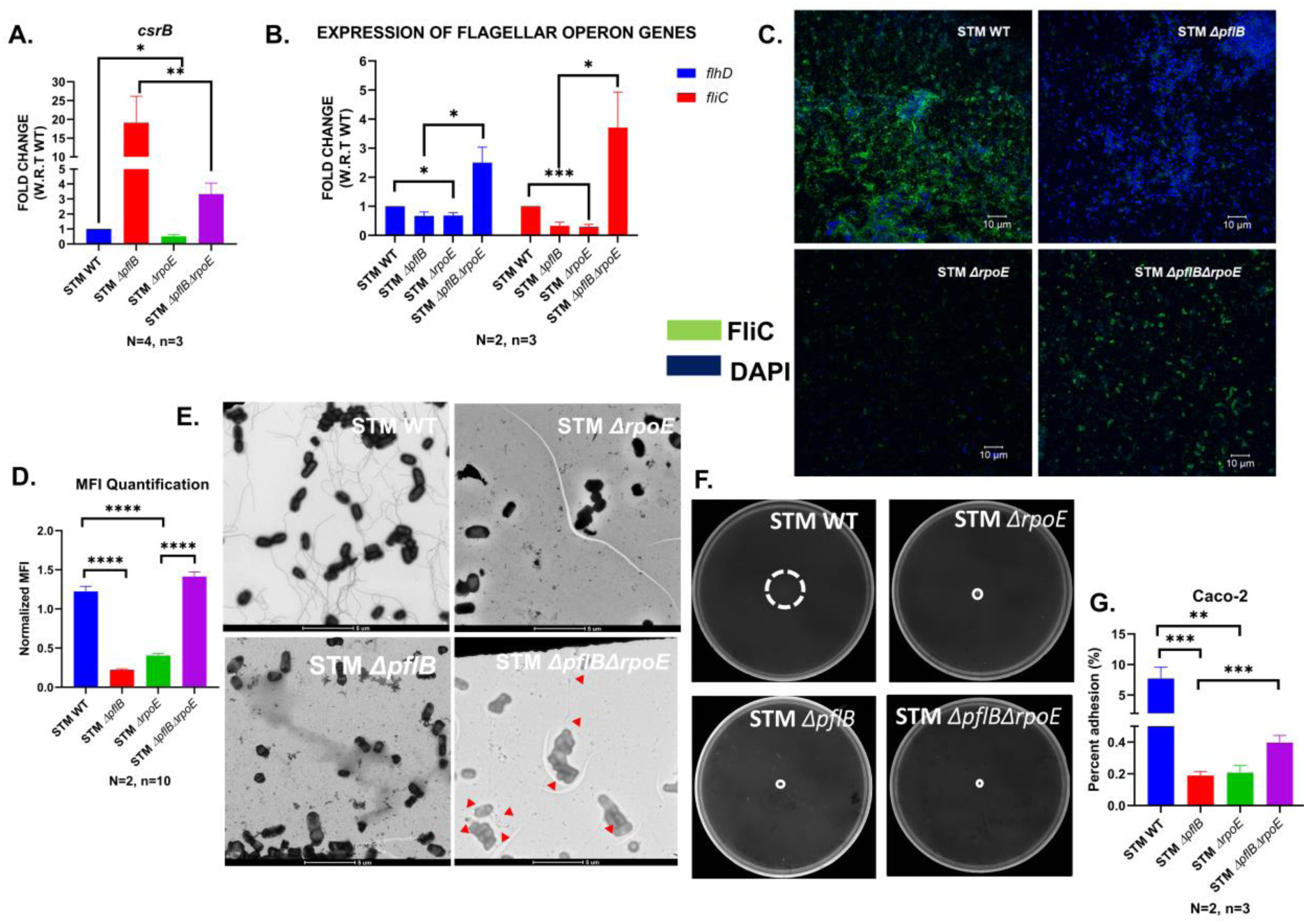
The extra cytoplasmic sigma factor RpoE increases the expression of *csrB* and therefore restricts the expression of *fliC*. A. RT-qPCR mediated expression profile of the sRNA *csrB* in the logarithmic cultures of STM WT, STM *ΔpflB*, STM *ΔrpoE*, STM *ΔpflBΔrpoE.* Data is represented as Mean+/- SEM of N=4, n=3. B. RT-qPCR mediated expression profile of the flagellar operon genes *flhD* and *fliC* in the logarithmic cultures of STM WT, STM *ΔpflB*, STM *ΔrpoE*, STM *ΔpflBΔrpoE*. Data is represented as Mean+/-SEM of N=2, n=3. C. Confocal microscopy assisted visualization of the flagellar structures in STM WT, STM *ΔpflB*, STM *ΔrpoE*, STM *ΔpflBΔrpoE*. Images are representative of N=2, n=10. D. Normalized Median Fluorescence Intensity (MFI), i.e.., MFI of Secondary antibody for Rabbit generated Anti-fli antibody/ MFI for DAPI in confocal microscopy assisted visualization of the flagellar structures in STM WT, STM *ΔpflB*, STM *ΔrpoE*, STM *ΔpflBΔrpoE*. Data is represented as Mean+/-SEM of N=2, n=10. E. TEM assisted visualization of the flagellar structures in STM WT, STM *ΔpflB*, STM *ΔrpoE*, STM *ΔpflBΔrpoE.* Data is representative of N=2, n=10. F. Swim motility of STM WT, STM *ΔpflB*, STM *ΔrpoE*, STM *ΔpflBΔrpoE* on 0.25% Agar containing LB plates G. Percent adhesion of STM WT, STM *ΔpflB*, STM *ΔrpoE*, STM *ΔpflBΔrpoE* in Caco-2 cell line. Data is represented as Mean+/-SEM of N=2, n=3. (Unpaired Student’s t-test for column graphs, Two-way Anova for grouped data, Mann-Whitney U-test for animal experiment data (**** p < 0.0001, *** p < 0.001, ** p<0.01, * p<0.05))

Our previous data demonstrated that formate produced by PflB helps maintain the intracellular pH (ipH) of the cell. Consequently, deletion of the *pflB* gene disrupts ipH balance, leading to membrane depolarization. We hypothesized that this membrane stress in *STM ΔpflB* could cause RseA degradation, thereby freeing σ^E^ to promote *csrB* transcription. Our results showed that the elevated *csrB* expression observed in *STM ΔpflB* was significantly reduced in *STM ΔpflBΔrpoE* (**Figure 5A**). Additionally, the suppressed expression of *flhD* and *fliC* in *STM ΔpflB* was restored in *STM ΔpflBΔrpoE* (**Figure 5B**). Confocal microscopy and TEM provided further confirmation of this finding (**Figure 5C-E**). However, we did not observe any recovery in the swim halo upon deletion of *rpoE* from STM *ΔpflB*, which shows that the recovery in translational level and protein level is not getting reflected in the motility of the attenuated strain (**Figure 5F**). However, there was an improved adhesion of *STM ΔpflBΔrpoE* to Caco-2 cells compared to *STM ΔpflB* (**Figure 5G**). This improved adhesion might be due to the enhanced hydrophobicity due to increased FliC protein in STM *ΔpflBΔrpoE* compared to STM *ΔpflB* [38]. These results suggest that the reduction in flagella synthesis in STM *ΔpflB* could be mediated by the extra cytoplasmic stress factor σ^E^.

### Formate regulates the balance between flagellar expression and SPI-1 gene expression in the intestine of C57BL/6 mice

Our previous data showed that *STM ΔpflB* exhibited a higher abundance of the *prgH* transcript. To determine if this increased expression of the SPI-1 gene was controlled by the same regulatory loop, we analysed the transcript levels of the SPI-1 genes *hilA* and *prgH*. We found that the elevated expression of *hilA* and *prgH* in *STM ΔpflB* was reduced in *STM ΔpflBΔrpoE* (**Figure 6A**). This confirmed that intracellular formate regulates the shift from flagellar expression to SPI-1 gene expression.

**Figure 6:**
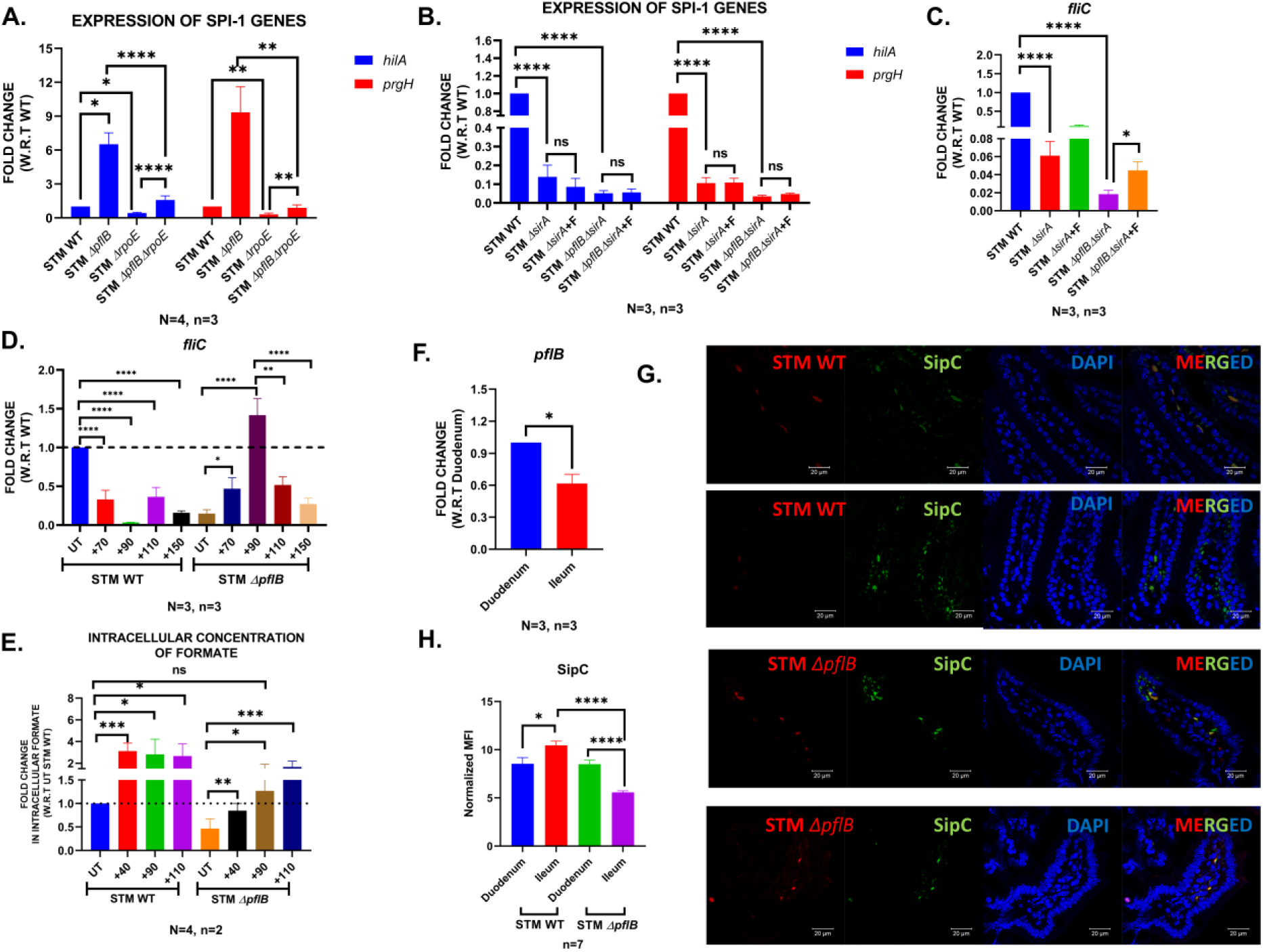
PflB maintains the switch between the flagellar expression and SPI-1 gene expression in intestine of the C57BL/6. A. RT-qPCR mediated expression profile of the SPI-1 genes *hilA* and *prgH* in the logarithmic cultures of STM WT, STM *ΔpflB*, STM *ΔrpoE*, STM *ΔpflBΔrpoE.* Data is represented as Mean+/-SEM of N=4, n=3. B. RT-qPCR mediated expression profile of the SPI-1 genes *hilA* and *prgH* in the logarithmic cultures of STM WT, STM *ΔpflB*, STM *ΔsirA*, STM *ΔpflBΔsirA.* Data is represented as Mean+/-SEM of N=4, n=3. C. RT-qPCR mediated expression profile of *fliC* gene in the logarithmic cultures of STM WT, STM *ΔpflB*, STM *ΔrpoE*, STM *ΔpflBΔrpoE.* Data is represented as Mean+/-SEM of N=3, n=3. D. RT-qPCR mediated expression profile of *fliC* gene in the logarithmic cultures of STM WT and STM *ΔpflB* supplemented with 70, 90, 110, 150 mM of sodium formate. Data is represented as Mean+/-SEM of N=3, n=3. E. GC-MS mediated quantification of relative intracellular formate in STM WT, STM *ΔpflB*, supplemented with 40, 90, 110 mM Sodium formate. Data is represented as Mean+/-SEM of N=4, n=2 F. RT-qPCR mediated expression profile of *pflB* gene in the duodenum and ileum of the intestine of C57BL/6 mice. Data is represented as Mean+/-SEM of N=3, n=3. G. Confocal microscopy assisted visualization of the SPI-1 effector SipC protein in the duodenum and ileum of mice infected with STM WT and STM *ΔpflB*. Images are representative of n=7. H. Confocal microscopy assisted visualization of the SPI-1 effector SipC protein in the duodenum and ileum of mice infected with STM WT and STM *ΔpflB*. Images are representative of n=7. (Unpaired Student’s t-test for column graphs, Two-way Anova for grouped data, Mann-Whitney U-test for animal experiment data (**** p < 0.0001, *** p < 0.001, ** p<0.01, * p<0.05))

Our previous results showed that formate supplementation enhances *prgH* expression in both STM WT and STM *ΔpflB* (**Figure 2A**). Flagellar motility and epithelial cell invasion are regulated by the BarA/SirA two-component system (TCS) [70]. As expected, STM *ΔsirA* exhibited low levels of *hilA* and *prgH* expression, and formate supplementation in STM *ΔsirA* did not significantly alter the expression of these genes. Additionally, deletion of *sirA* in STM *ΔpflB* eliminated any significant upregulation of *hilA* and *prgH* in response to formate supplementation (**Figure 6B**). This indicates that extracellular formate influences the virulence and pathogenicity of *Salmonella* via the BarA/SirA TCS.

Interestingly, we observed that STM *ΔpflBΔsirA* continued to show recovery in *fliC* transcript levels under formate supplementation. This suggests that intracellular formate regulates *Salmonella* pathogenesis independently of the BarA/SirA TCS (**Figure 6C**). Therefore, when the intracellular formate pool is maintained by the *pflB* gene, extracellular formate modulates SPI-1 gene expression through the BarA/SirA TCS. However, if the intracellular formate pool is disrupted, the resulting membrane stress triggers RpoE-mediated overexpression of *csrB*, which is associated with reduced *fliC* transcript levels and increased *hilA* and *prgH* expression.

As expected, STM *ΔsirA* exhibited low expression levels of *hilA* and *prgH*, and formate (F) supplementation in STM *ΔsirA* did not result in any significant change in the expression of these genes. Furthermore, deletion of *sirA* in STM *ΔpflB* eliminated any significant upregulation of *hilA* and *prgH* in response to formate supplementation (**Figure 6B**). This demonstrates that extracellular formate influences the virulence and pathogenesis of *Salmonella* through the BarA/SirA TCS. Interestingly, we observed that STM *ΔpflBΔsirA* continued to show recovery in *fliC* transcript levels with formate supplementation. This indicates that intracellular formate regulates *Salmonella* pathogenesis independently of the BarA/SirA TCS (**Figure 6C**). Thus, when the intracellular formate pool is maintained by the *pflB* gene, extracellular formate modulates SPI-1 gene expression via the BarA/SirA TCS. However, if the intracellular formate pool is disrupted, the resulting membrane stress triggers RpoE-mediated overexpression of *csrB*, which is associated with reduced *fliC* transcript levels and increased *hilA* and *prgH* expression. We proceeded to understand whether achieving intracellular formate concentrations of STM *ΔpflB* to similar levels as STM WT and further supplementing it with formate can replicate the behaviour of wild-type strain in presence of formate. We subcultured the stationary phase cultures of STM WT and STM *ΔpflB* and supplemented them higher concentrations of formate (70, 90, 110, 150 mM). We found that there was a substantial decrease in the in the *fliC* transcript level in STM WT upon supplementation of higher concentrations of formate. STM *ΔpflB* showed a remarkable recovery in the *fliC* transcript level at 90 mM concentration, following which the expression levels declined in the presence of 110 mM and 150 mM formate (**Figure 6D**). GC-MS mediated quantification of intracellular formate concentration revealed that a concentration of 90 mM was enough to replenish the depleted intracellular formate pool in STM *ΔpflB,* which explains the highest expression of *fliC* at this concentration (**Figure 6E**). Beyond this, supplementation of formate reduces the expression of *fliC* as seen in **Figure 6D**. This mimics the scenario of supplementing F to STM WT with an intact formate pool. This was also visualized under TEM and quantified by confocal microscopy (**Figure S6A, B, C**).

To explore the physiological relevance of this phenomenon, we examined *pflB* expression levels in different sections of the intestine, since flagella and SPI-1 play critical roles in the intestinal phase of *Salmonella* infection. We observed that *pflB* expression decreased progressively from the duodenum to the ileum (**Figure 6F**). This finding led us to hypothesize that *Salmonella* downregulates its *pflB* gene as it approaches its invasion site—the distal ileum.

This downregulation could induce a partial formate deficiency within the bacterial cell, causing membrane depolarization, which in turn activates σ^E^, increases *csrB* abundance, downregulates flagellar operon transcription, and upregulates the SPI-1 gene cascade, ultimately help in bacterial invasion at its target site of ileum.

To test this hypothesis, we collected duodenum and ileum sections from mice infected with STM WT and STM *ΔpflB* and stained them with an anti-SipC antibody to measure levels of the SPI-1 effector protein (**Figure 6G**). We normalized the fluorescence from the secondary antibody, specific to the rabbit anti-SipC antibody, against mCherry fluorescence from bacteria transformed with the pFPV plasmid. In mice infected with STM WT, we found that the normalized mean fluorescence intensity (MFI) increased from the duodenum to the ileum. In contrast, STM *ΔpflB*-infected mice showed a decrease in normalized MFI from the duodenum to the ileum (**Figure 6H**). These results indicate that the formate pool maintained by the PflB enzyme regulates SPI-1 gene expression, and its depletion may be a strategy used by *Salmonella* Typhimurium to maximize SPI-1 expression, and thereby invasion, in the ileum.

## Discussion

*Salmonella* possesses a network of genes and regulatory factors that manages the balance between its extracellular and intracellular modes of existence [82]. Research on *Salmonella* Typhimurium has demonstrated overlapping regulatory mechanisms between the flagellar secretion system and the SPI-1. The flagellar protein FliZ is known to activate the primary regulator of flagella synthesis, FlhD, through post-translational modification. Kage *et al*. have shown that FliZ can also regulate HilD, a key regulator of SPI-1, at the transcriptional level [83, 84]. HilD has been found to bind directly to the promoter region of *flhDC*, upstream of the P5 transcriptional start site, thus enhancing its expression [85]. Other studies indicate that SPI-1 expression can suppress flagellar gene expression, aiding in pathogenicity and virulence of *Salmonella*. A mutation in *fliZ* reduces *hilA* expression by half, while a knockout of *sirA* decreases *hilA* expression tenfold and increases flagella expression a hundredfold [86-89]. Furthermore, recent research by Saleh *et al*. has shown that HilD activation can trigger a SPI-1 dependent induction of the stringent response and significantly reduce the Proton Motive Force (PMF). Although flagellation remains unaffected, HilD activation results in a motility defect in *S.* Typhimurium [90]. Collectively, these studies reveal that *Salmonella* can toggle between flagella synthesis and SPI-1 gene expression to optimize adhesion and the delivery of SPI-1 effectors. There are several potential benefits to downregulating flagella while upregulating SPI-1 genes. First, reducing flagella expression prevents the bacteria from moving away from a target cell primed for invasion. Second, because the flagellar apparatus is immunogenic, downregulating it may help the bacteria evade the host immune response [91-93]. Lastly, simultaneous production of both systems could lead to interference, potentially causing the improper secretion of flagellin and SPI-1 effectors [94].

In our study, we knocked out the gene encoding the pyruvate-formate lyase enzyme, effectively depleting the cell of its native formate pool. Although the knockout strain displayed increased expression of the SPI-1 gene *prgH*, it showed reduced transcript levels of *fliC*. This was reflected in decreased invasion and adhesion capabilities in a human epithelial cell line. These deficiencies were partially restored with the addition of 40 mM formate, which also explains the strain’s hyperproliferation in the RAW 264.7 macrophage cell line [95]. Additionally, we found that the STM Δ*pflB* mutant was unable to maintain its intracellular pH to the same extent as the STM wild type, resulting in a membrane depolarization phenotype. Membrane stress triggered the sigma factor RpoE to increase the expression of *csrB*. Since *csrB* is known to bind to and inhibit the CsrA protein, we hypothesized that the elevated *csrB* transcript levels led to reduced *fliC* expression. Overexpressing *csrA* via an extragenous plasmid partially restored *fliC* expression, supporting our hypothesis. Furthermore, deletion of *rpoE* in the STM Δ*pflB* strain successfully reversed the flagella-deficient phenotype, providing further confirmation. This demonstrated that the extra-cytoplasmic sigma factor RpoE mediates downstream effects in response to cell membrane stress.

In all our experiments, we also observed that formate supplementation led to reduced flagellar expression in STM WT. Interestingly, adding formate did not affect the intracellular pH or membrane potential of STM WT, which led us to hypothesize that extracellular formate impacts STM pathogenicity differently than intracellular formate synthesis. Huang *et al*. demonstrated that extracellular formate functions as a diffusible signal to enhance invasion, but this effect was independent of the BarA/SirA Two-Component System and was also unrelated to *csrB*, which is directly regulated by the response regulator SirA. In our study, we found that STM WT with formate supplementation (STM WT+F) showed higher expression of SPI-1 genes [96]. Contrary to previous findings, we observed that *csrB* transcript levels were higher in STM WT+F compared to STM WT, suggesting that elevated *csrB* levels might be linked to reduced *fliC* expression [97]. This indicates that formate is a crucial metabolite for *Salmonella*, with a critical concentration determining whether it acts through the TCS pathway or by regulating the cell’s intracellular pH. Additionally, we found that supplementing extracellular formate to restore the intracellular formate pool caused STM Δ*pflB* to mimic the wild-type strain, exhibiting reduced *fliC* levels in the presence of formate.

*Salmonella* Typhimurium has been extensively studied for its ability to adapt its metabolism to conditions within the host. Previous research has shown that spatially distinct *Salmonella* populations in the mucous layer and intestinal lumen differ in their energy metabolism. Transcript levels of pyruvate-formate lyase and formate dehydrogenases were found to be higher in the mucous layer than in the lumen population [16]. This formate metabolism provided a fitness advantage in the mucous layer. In our study, we observed that *pflB* transcription was highest in the large intestine and gradually decreased in the small intestine, the primary site of *Salmonella* infection. This suggests that *Salmonella* modulates *pflB* expression in response to local gut microenvironment signals. We propose a model in which *Salmonella* downregulates the *pflB* gene upon reaching its infection site, disrupting its intracellular formate pool. This can result in partial membrane depolarization, which downregulates the flagellar apparatus and promotes invasion into epithelial cells through upregulated expression of the SPI-1 system. The specific signals and genetic mechanisms controlling this modulation remain unclear.

Overall, our study contributes to understanding how intracellular metabolism in *Salmonella* influences its ability to adhere to and invade non-phagocytic intestinal epithelial cells, and it highlights the potential for uncovering additional host-pathogen metabolic interactions in future research.

## Acknowledgements

We acknowledge the Departmental Real-Time Facility, Departmental Confocal Microscopy Facility, Divisional Electron Microscopy Facility, Divisional Mass Spectrometry Facility, Divisional Flow Cytometry Facility, and Central Animal Facility at IISc for providing us opportunity for experimentation. Mr. Sumitlal K and Ms. Navya have helped in image acquisition in Confocal Microscopy. Dr. Sushma and Keerthana helped with image acquisition in Transmission Electron Microscopy. Dr. Muralidhar and Dr. Sunitha helped in formate quantitation studies using Gas Chromatography Mass Spectrometry. Ms. Lipika, Ms. Dharani, and Mr. Munish helped in data acquisition from Flow Cytometry. Dr. Akshay Datey is acknowledged for his initial supervision in the study. The anti-SipC antibody was a kind gift from Prof. Dr. Ayub Qadri, National Institute of Immunology, Delhi, India.

## Author Contributions

DM and DC have conceived the study and designed the experiments. DM has performed all the experiments and participated in acquiring the data. DM has analysed the data, constructed the figures, and wrote the original draft of the manuscript. DM and DC have participated in the proofreading and editing of the manuscript. DC has supervised the study and helped in fund acquisition. All the authors have read and approved the manuscript.

## Data availability

Data can be available from the authors upon reasonable request.

## Data deposition

Link for pre-print: https://doi.org/10.1101/2024.01.30.577997

## Thesis link

This research has not yet been included as part of any PhD thesis till current date.

---

**Supplementary Figure 1:**
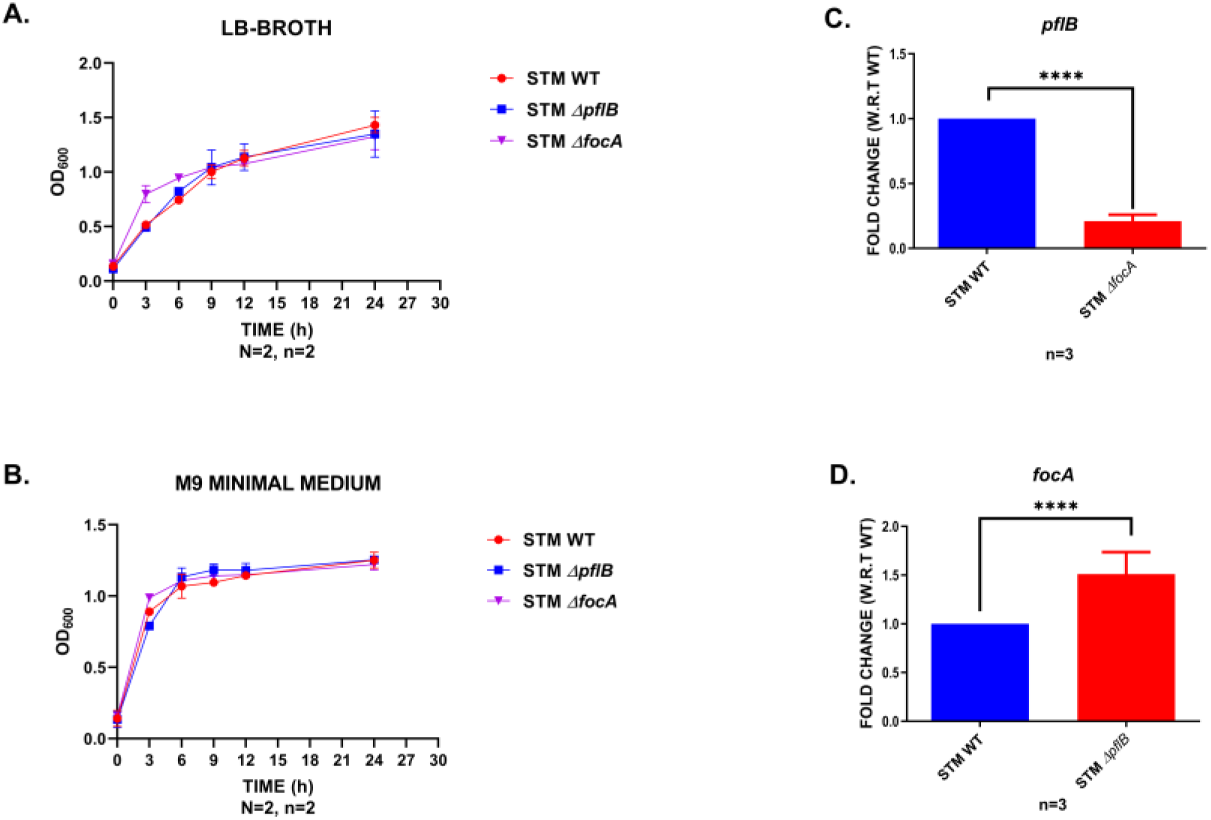
STM *ΔpflB*, and STM *ΔfocA* are not growth deficient in LB Broth, M9 Minimal Media. Deletion of pflB caused an enhanced expression of *focA*. A. Growth curves of STM WT, STM *ΔpflB* and STM *ΔfocA* in LB broth, M9 Minimal Media. Data is represented as Mean+/-SEM of N=2, n=2. B. Growth curves of STM WT, STM *ΔpflB* and STM *ΔfocA* in M9 Minimal Media. Data is represented as Mean+/-SEM of N=2, n=2. C. RT-qPCR mediated expression profile of *pflB* gene in the logarithmic cultures of STM *ΔfocA*. Data is represented as Mean+/-SD of n=3. D. RT-qPCR mediated expression profile of *focA* gene in the logarithmic cultures of STM *ΔpflB*. Data is represented as Mean+/-SD of n=3. (Unpaired Student’s t-test for column graphs, Two-way Anova for grouped data, Mann-Whitney U-test for animal experiment data (**** p < 0.0001, *** p < 0.001, ** p<0.01, * p<0.05))

**Supplementary Figure 2:**
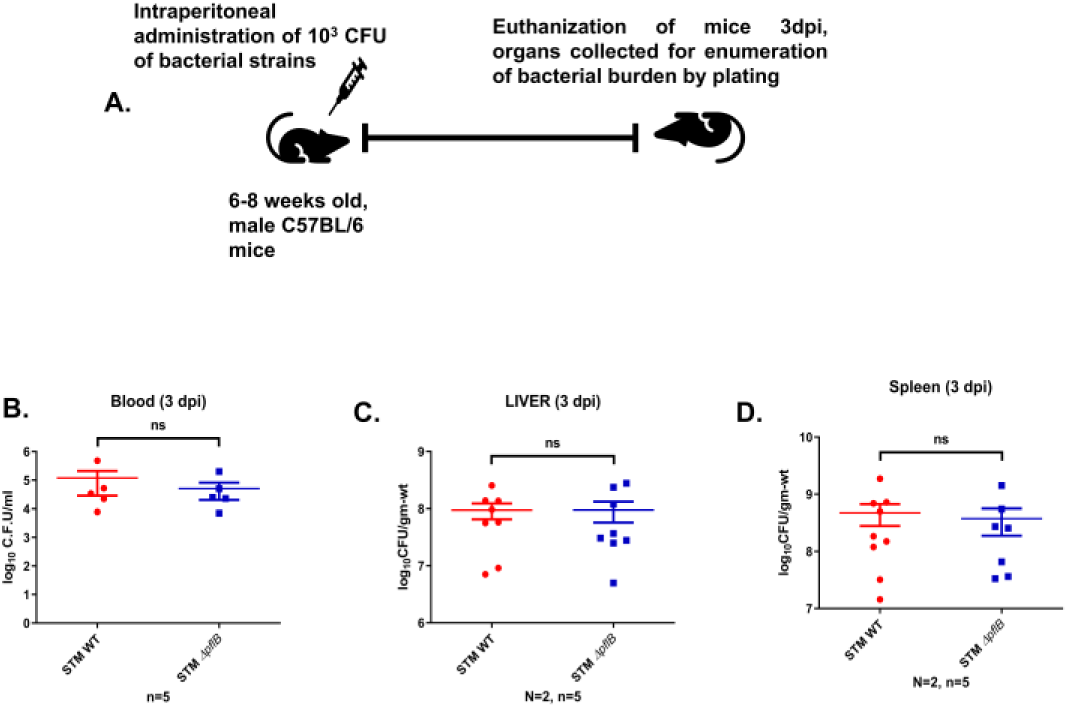
Intraperitoneal infection of STM WT and STM *ΔpflB* lead to no significant change in the bacterial burden in blood, liver, and spleen 3 dpi. A. Schematic showing the protocol followed for determining the organ burden of STM WT and STM *ΔpflB* 3 days post intraperitoneal infection B-D. Bacterial burden of STM WT and STM *ΔpflB* in blood (B), liver (C), and spleen. Data is represented as Mean+/-SEM of N=2, n=5. (Unpaired Student’s t-test for column graphs, Two-way Anova for grouped data, Mann-Whitney U-test for animal experiment data (**** p < 0.0001, *** p < 0.001, ** p<0.01, * p<0.05))

**Supplementary Figure 3:**
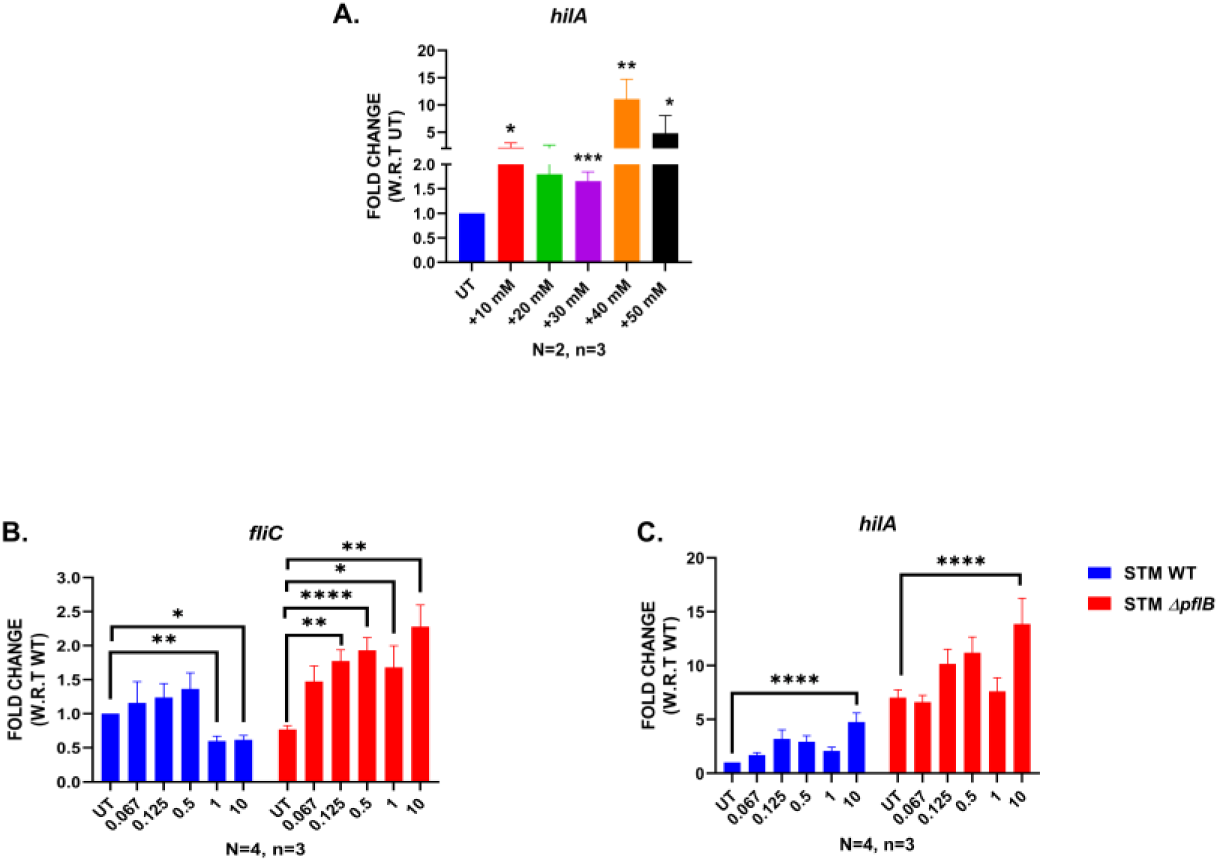
Formate modulates the expression of *fliC* and *hilA* in STM WT and STM *ΔpflB* in a concentration dependent manner. A. Expression of *hilA* in STM WT in formate supplemented at concentrations of 10, 20, 30, 40, and 50 mM. Data is representative of N=2, n=3 and expressed as Mean+/- SD. B. Expression of *fliC* in STM WT and STM *ΔpflB* in formate supplemented at concentrations of 0.067, 0.125, 0.5, 1, 10 mM. Data is represented as Mean+/-SEM of N=4, n=3. C. Expression of *hilA* in STM WT and STM *ΔpflB* in formate supplemented at concentrations of 0.067, 0.125, 0.5, 1, 10 mM. Data is represented as Mean+/-SEM of N=4, n=3. (Unpaired Student’s t-test for column graphs, Two-way Anova for grouped data, Mann-Whitney U-test for animal experiment data (**** p < 0.0001, *** p < 0.001, ** p<0.01, * p<0.05))

**Supplementary Figure 4:**
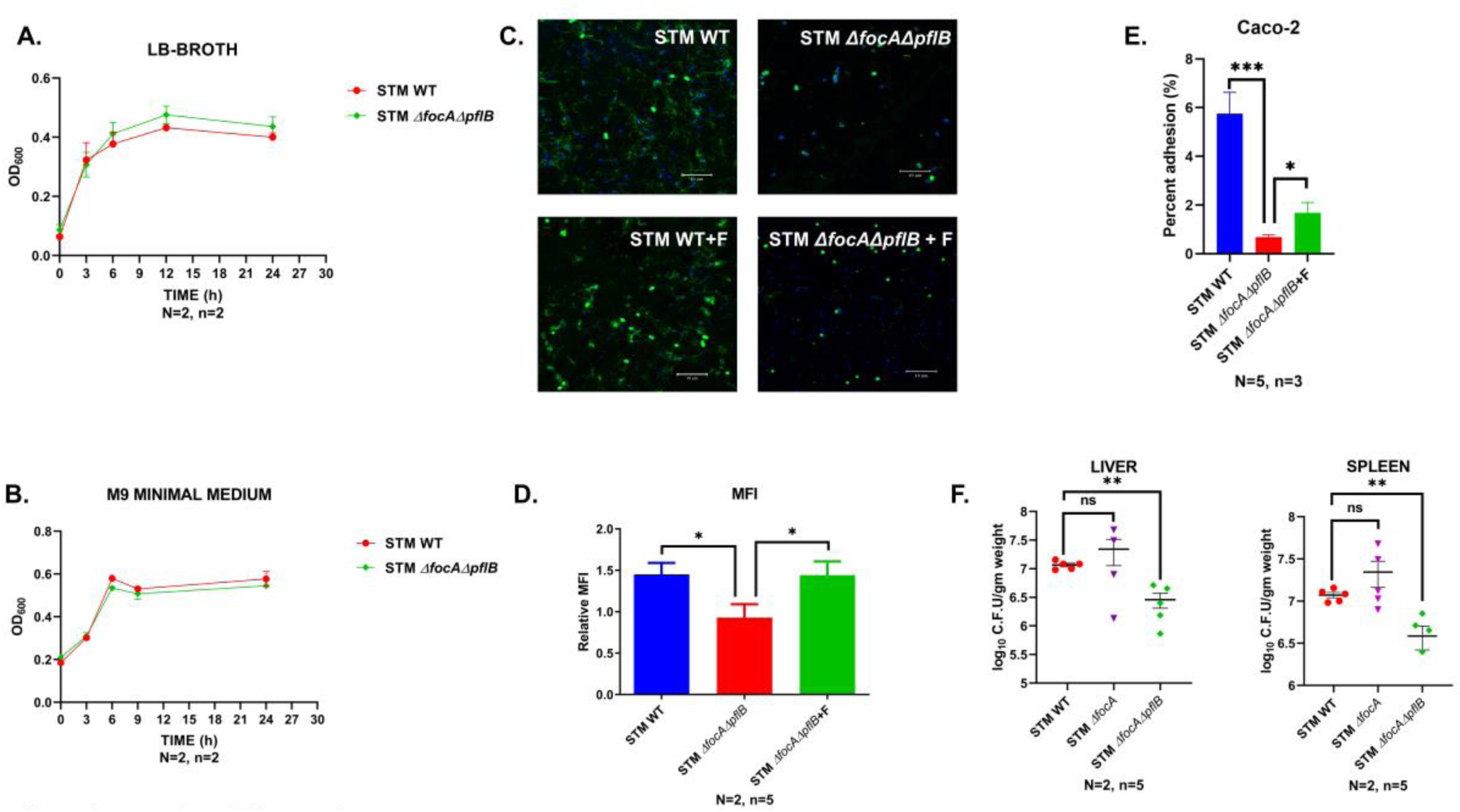
STM *ΔfocAΔpflB* had an attenuated flagellar apparatus, with partial recovery upon supplementation of extragenous F. A. Growth curves of STM WT, STM *ΔfocAΔpflB* in LB broth and M9 Minimal Media. Data is represented as Mean+/-SEM of N=2, n=2. B. Growth curves of STM WT, STM *ΔpflB* and STM *ΔfocA* in M9 Minimal Media. Data is represented as Mean+/-SEM of N=2, n=2. C. Confocal microscopy assisted visualization of the flagellar structures in STM WT, STM *ΔfocAΔpflB* (+/-F). Images are representative of N=2, n=5. D. Normalized Median Fluorescence Intensity (MFI), i.e.., MFI of Secondary antibody for Rabbit generated Anti-fli antibody/ MFI for DAPI in confocal microscopy assisted visualization of the flagellar structures in STM WT, STM *ΔfocAΔpflB* (+/-F). Data is represented as Mean+/-SEM of N=2, n=5. E. Percent adhesion of STM WT and STM *ΔfocAΔpflB* in Caco-2 cell line. Data is represented as Mean+/-SEM of N=5, n=3. F. Organ burden of STM WT and STM *ΔfocAΔpflB* in liver and spleen of C57BL/6 mice 5 days post oral gavaging. Data is represented as Mean+/-SEM of N=2, n=5. (Unpaired Student’s t-test for column graphs, Two-way Anova for grouped data, Mann-Whitney U-test for animal experiment data (**** p < 0.0001, *** p < 0.001, ** p<0.01, * p<0.05))

**Supplementary Figure 5:**
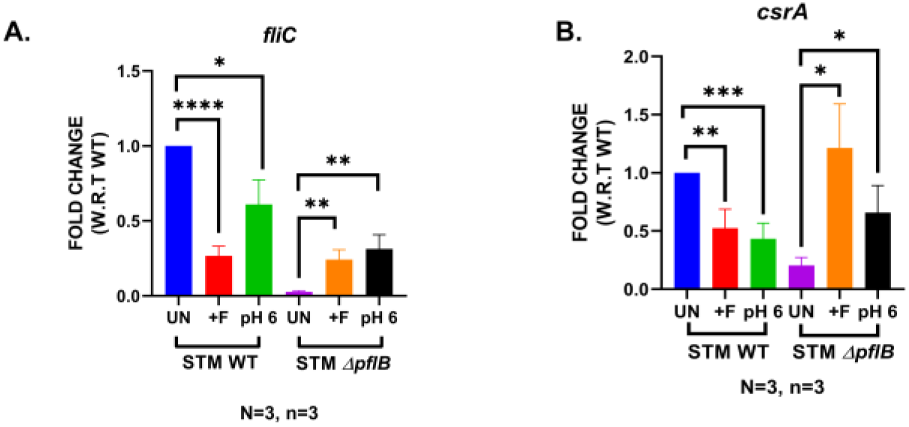
Decreased expression of *fliC* in STM *ΔpflB* can be altered upon incubating the bacterial cultures in acidic media (pH of 6). **A.** RT-qPCR mediated expression profile of *fliC* in the logarithmic cultures of STM WT, STM *ΔpflB* incubated in acidic LB media (pH 6). Data is represented as Mean+/-SEM of N=3, n=3. **B.** RT-qPCR mediated expression profile of *csrA* in the logarithmic cultures of STM WT, STM *ΔpflB* incubated in acidic LB media (pH 6). Data is represented as Mean+/-SEM of N=3, n=3. (Unpaired Student’s t-test for column graphs, Two-way Anova for grouped data, Mann-Whitney U-test for animal experiment data (**** p < 0.0001, *** p < 0.001, ** p<0.01, * p<0.05))

**Supplementary Figure 6:**
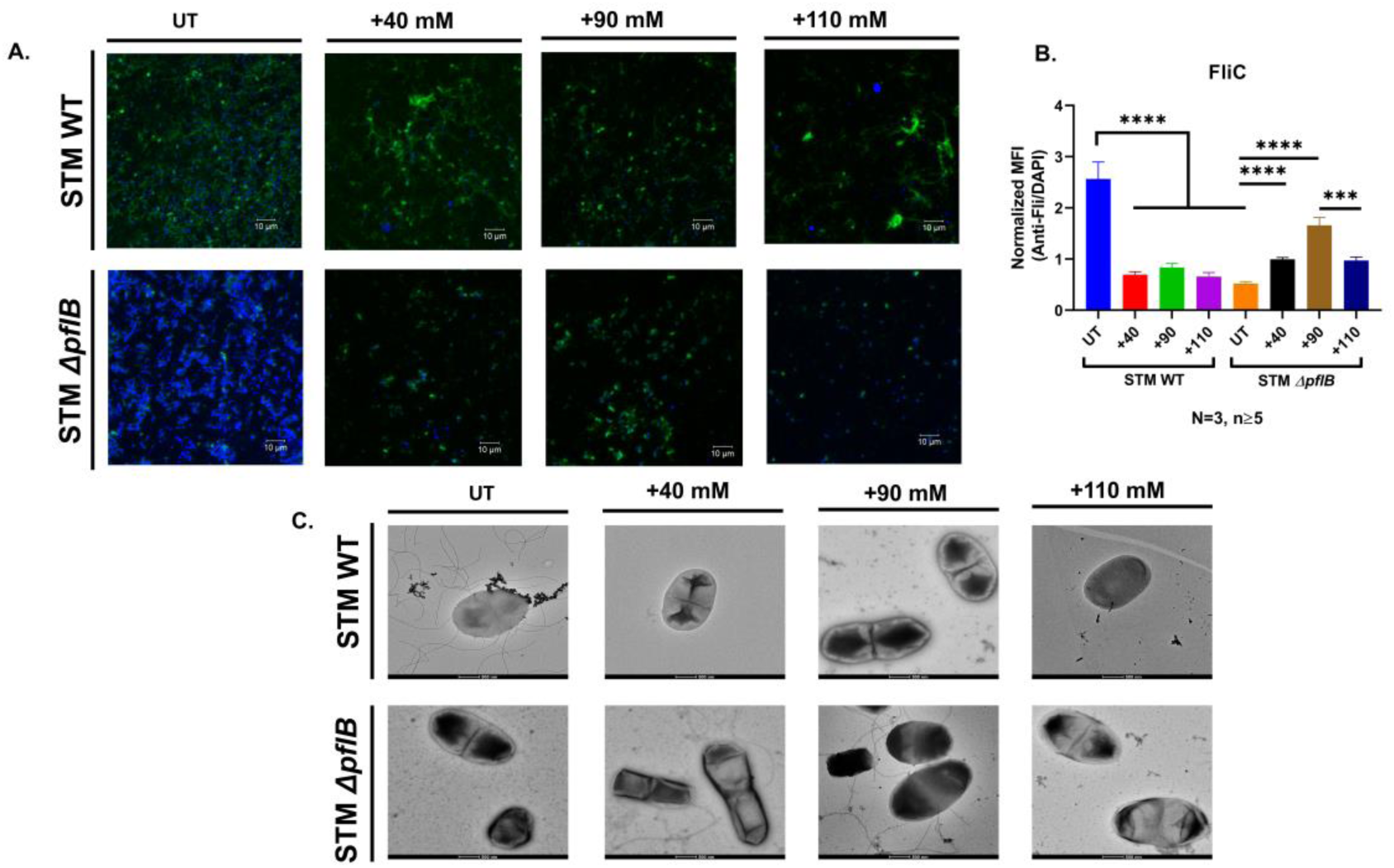
Compensation of intracellular formate pool in STM *ΔpflB* makes it mimic the behaviour of STM WT. A. Confocal microscopy assisted visualization of the flagellar structures in STM WT, STM *ΔpflB* supplemented with 40, 90, 110 mM of sodium formate. Images are representative of N=3, n≥5. B. Normalized Median Fluorescence Intensity (MFI), i.e.., MFI of Secondary antibody for Rabbit generated Anti-fli antibody/ MFI for DAPI in confocal microscopy assisted visualization of the flagellar structures in STM WT, STM *ΔpflB* supplemented with 40, 90, 110 mM of sodium formate. Data is represented as Mean+/-SEM of N=3, n≥5. C. TEM assisted visualization of the flagellar structures in STM WT and STM *ΔpflB* supplemented with 40, 90, 110 mM of sodium formate. Data is representative of N=2, n=10.

